# Clustering of Inverted Triplications in Centromeric and Subtelomeric Chromosomal Regions of *Aspergillus flavus*

**DOI:** 10.1101/2025.01.14.632981

**Authors:** Jeffrey W. Cary, Amna Malik, Alexie Smith, Rose Ludecke, Erik Flemington, Arthur J. Lustig

## Abstract

The formation of inverted repeats frequently initiates rearrangements associated with gene amplification. Among these rearrangements are inverted triplications (TRP/INVs) that are associated with multiple disease states in humans. We have used the filamentous fungus Aspergillus flavus as a model system to analyze spontaneous DNA rearrangements using high-depth third-generation sequencing of DNA isolated from vegetatively cultured strains. Analysis of sequence data identified a class of infrequent and transient rearrangements having structures typical of TRP/INVs. These TRP/INVs form both unprocessed and processed species and have a variably sized deletion at the junction between direct and inverted sequences. We found that TRP/INVs are enriched in heterochromatic centromeric and subtelomeric A+T-rich regions, suggesting that this process is a source of genetic instability in these domains. Consistent with this finding, A+T-rich regions contain elevated levels of direct, inverted, and perfect palindromic repeats. Inverted junctions contain palindromic sequences that are typically associated with TRP/INVs. A closer examination of the class of genomic palindromes found at inverted junctions revealed a broad distribution, with the majority lying in AT-rich centromeric sequences. The distribution of TRP/INVs in centromeric and subtelomeric domains mirrors the frequency of palindromes, indicating that palindrome abundance is likely to be a driver of TRP/INVs. A further examination of predicted pairing patterns in palindrome-like structures suggests that these structures may function as sites of DNA resolution, strand transfer, and replicative stalling. These results are consistent with a replication-based model in which palindromes, under conditions of replication stress, facilitate the formation of TRP/INV structures. These studies also represent, to our knowledge, the first characterization of spontaneous TRP/INV formation, likely to underlie the complex rearrangements associated with higher eukaryotic genomic instability and disease states.

**AUTHOR SUMMARY:** Inverted triplications (TRP/INVs), coupled with gene amplification, are sources of genetic instability. However, little is known about how these species arise spontaneously prior to selective pressures. We have established a fungal model system that uses long-read third-generation sequencing to identify low-abundance rearrangements in the absence of an intentional selection. We identified and characterized multiple classes of TRP/INVs flanked by palindromic sequences. These rearrangements were concentrated in the AT-rich centromeric and subtelomeric regions, suggesting that these sites are regulated recombinational hotspots. Palindromes that are also associated with TRP/INVs are also most prevalent in A+T-rich centromeric regions, suggesting that the abundance of palindromes, both as a source of DNA damage and fork stalling, is a driver of TRP/INV formation. These studies provide the first evidence (to our knowledge) of the spontaneous formation of TRP/INVs during DNA replication, which is likely to precede more complex sources of rearrangements associated with oncogenesis and other disease states in higher eukaryotes.

## INTRODUCTION

Gene duplication is a common theme in genetic change in all phyla ranging from bacterial to fungal to human cells and has been implicated in the genesis of disease states [1, 2]. Indeed, the spontaneous formation of large duplication inversions is an underlying mechanism contributing to oncogenic states [2, 3].

Model systems are valuable tools to select and characterize gene duplications. Duplications can occur with long sequence elements through transposition and homologous recombination. Rearrangements, however, also occur in the presence of minimal homology, particularly in the presence of perfect paired and imperfect (quasi-) palindromes [4]. Palindromes are particularly unstable due to their extrusion as either cruciform or hairpin structures. For example, breakage-fusion-bridge (BFB) cycles can involve either the loss of a telomere, or, alternatively the processing of palindromic cruciform DNA to form inversions and subsequent DNA breakage, followed by fusion of DNA strands through processes such as non-homologous end joining [3, 5–7].

Inverted triplications (TRP/INVs) are another class of species mediated by the rearrangement of relatively short regions of homology [8–11]. These latter species are observed in bacteria and yeast and can be initiated by palindromes under conditions of replicative stress and fork regression during bi-directional DNA replication [12, 13]. Recent data has also suggested a role for TRP/INVs in the rearrangements involved in a multiplicity of disease states [14]. In one model for the formation of these structures, palindromes flanking a gene can create a hairpin that primes DNA replication and the synthesis of the inverted copy. After reaching a second palindrome, template switching and DNA replication give rise to the direct repeat. The junctions between direct and inverted repeats are thought to reform the palindromes in the resulting TRP/INV, which may be resolved into asymmetric TRP/INVs containing deletions surrounding the palindromic junctional sequence. On a statistical basis, short palindromes are expected to form most easily in AT or GC-rich sequences. AT-rich palindromes form frequently in the human genome and are involved in both chromosomal translocation and evolutionary hypervariability [15, 16]. TRP/INVs and other rearrangements are typically observed only under selection. However, the array of rearrangements that may occur in the absence of any selection or in regions of simple sequence non-genic DNA have not been investigated.

Centromeric and telomeric regions of chromosomes are essential for the stable inheritance of genetic information. At the microtubule attachment site, centromeric domains can be present as point or regional structures, the latter including chromosome-specific satellite sequences [17]. In *Neurospora* and other filamentous fungi, centromeres appear to be embedded within heterochromatic A+T-rich sequences, modeled in part by a centromere-specific histone H3 variant [18, 19]. These centromeres are thought to be generated through a process of repeat-induced point mutation (RIP) [20–22]. In another filamentous fungus, *Aspergillus (A.) flavus*, centromeres consist of approximately 100 kb of 90-95% % A+T rich sequence on each of the eight chromosomes [23, 24]. Generally, centromere sequences on homologs in different strains share extensive (88-99%), but not complete, homology [24]. In contrast, the AT-rich sequences of non-homologs share minimal homology. Thus, competing forces appear to maintain a balance between variation and conservation.

Rearrangements near fungal telomeres are thought to drive genetic reshuffling, adaptive polymorphisms, and elimination of gene sequences [23, 25]. The subtelomeric 100 kb, adjacent to the telomeric (TTAGGGTCAACA)_9-11_ repeats [26], is also interspersed with A+T-rich elements (ATEs), of 2-20 kb in length, that are distributed in different configurations among the chromosomal arms within and between strains. Our previous studies have revealed remnant transposable elements in subtelomeres, suggesting that repetitive cycles of transposition and RIP contribute to the evolution of these sequences [24]. Nonetheless, we found highly conserved ATEs that lack any homology to transposable elements on both the same chromosome of different strains and distinct chromosomes of the same strain. Hence, subtelomeric ATEs are also maintained in a homeostasis of divergence and conservation, indicating a process that acts to promote recombination. The processes acting to promote and to limit ATE destabilization are not understood.

Identifying spontaneous rearrangements using standard genetic techniques has been challenging, due to their low frequencies and the limited means to identify complex forms [27]. We have employed a combination of two approaches to examine species arising from spontaneous events. First, the multinucleate mycelial growth of *A. flavus* leads to the potential identification of rare deleterious mutations and rearrangements [28]. Selection would tend to eliminate such species. Second, we used analysis of high-coverage third-generation sequencing as a physical readout of these genetic rearrangements. Here, we report on the identification of infrequent and transient TRP/INVs clustered in heterochromatic centromeric and subtelomeric regions in the absence of overt selection; a process that appears to be driven in part by the abundance of small palindromes.

## RESULTS

### A+T Rich Elements in Centromeres and Subtelomeric Regions Have a Nonrandom Distribution of Direct, Inverted, and Palindromic Repeats

To ascertain the accumulation of short repeated sequences in *A. flavus* AT-rich centromeric regions, we examined random 250 bp DNAs for BLAST self-homology and identified potential direct, inverted, and palindromic repeats. We plotted the BLAST self-homology e-values [29] of each repeat class in 129 randomly selected 250 bp, 90% A+T-rich sequences from centromeric regions. This result was then compared to repeats identified among 500 computationally derived and random 250 bp, 90% A+T-rich sequences (Table 1).

**Table 1.** Direct, Inverted and Palindromic Repeats in Centromeric, Subtelomeric and Randomly Generated A+T Rich tracts. The sequence source, repeat type, BLAST e-value, and number of samples assayed containing either ≥ 1 or 0 repeat are listed together with the p values from a comparison of the genomic and random sequences as determined by the Fisher Exact Test (FET).

| Sequence | Repeat | BLAST e value | $\geq 1$ Repeats | 0 Repeats | P value (FET) |
| --- | --- | --- | --- | --- | --- |
| Centromeric | Direct | $<10^{-4}$ | 13 | 116 | $<0.0001$ |
| Subtelomeric | Direct | $<10^{-4}$ | 5 | 75 | $<0.02$ |
| Random | Direct | $<10^{-4}$ | 7 | 493 | NA |
| Centromeric | Inverted | $<10^{-4}$ | 9 | 120 | $<0.001$ |
| Subtelomeric | Inverted | $<10^{-4}$ | 6 | 74 | $<0.003$ |
| Random | Inverted | $<10^{-4}$ | 6 | 494 | NA |
| Centromeric | Palindrome | $<10^{-8}$ | 9 | 120 | $<0.003$ |
| Subtelomeric | Palindrome | $<10^{-8}$ | 6 | 74 | $<0.007$ |
| Random | Palindrome | $<10^{-8}$ | 8 | 492 | NA |

The most significant inverted and direct repeats (e-values <10^-4^) identified in either genomic or randomly generated sequences were compared by the Fischer Exact Test. We determined that the direct repeats within centromeres were at significantly higher values than those generated from random sequences (p<0.0001). We further identified a significantly higher level of inverted repeats (p<0.001). Due to the higher spontaneous level of palindromic sequences in AT-rich sequences, we compared centromeric and random sequences at BLAST e-values of <10^-8^ and found a significant increase in the number of perfect palindromes (p<0.003) in centromeres. Palindromic sequences were found in 6.9% of centromeres sequences that were tested.

We conducted an analogous test among eighty, 250 bp sequences extracted from ATEs within subtelomeric regions, defined as the 100 kb adjacent to the telomeric (TTAGGGTCAACA)_9-11_ repeats [23, 25] (Table 1). Direct and inverted repeats were quantified in this region and compared with randomized sequences. Both sets of repeats with e values <10^-4^ were present at elevated levels in the subtelomere, relative to random sequences (direct repeats: p<0.002; inverted repeats: p<0.002). Similarly, perfect palindromic sequences were again present more frequently in subtelomeric sequence than in random sequence datasets (p<0.003) and were present in 7.5% of the subtelomeric sequences. Hence, all three classes of repeats were significantly elevated in centromeric and subtelomeric domains, suggesting an active process that generates this nonrandom behavior.

### Qualitative Identification of a TRP/INV Class in Cultured Cells

We were interested in identifying spontaneous rearrangements that may help to explain the nonrandom characteristic of repeated elements in subtelomeric and centromeric ATE sequences. To this end, we carried out high coverage (500-1500-fold) third-generation Oxford nanopore (ONT) sequencing as a readout for infrequent structural variants (SVs) [30] after forced vegetative culturing. We optimized the Sniffles 2 and minimap2 programs [31–33] for infrequent SVs and categorized them into apparent deletions, insertions, inversions, duplications, and translocations. We specifically characterized individual sequencing reads from the insertion class that predict sizes of ≥ 300 bp. BLAST analysis of these larger insertion sequences revealed two major classes: intact tandem duplications and duplications interrupted by a partial or complete inversion. This study focused on the latter class of structures that have structures akin to the inverted triplications [TRP/INVs], previously described in gene amplification studies. [8–10]

Our initial qualitative analysis compared this subset of the SV insertion class that appeared to arise in clusters from *A. flavus* genomic DNA prepared from a starter culture (G0) (generated from spores isolated from conidiophores) after five (G5), ten (G10), and fifteen (G15) weeks of forced vegetative (mycelial) growth. One of the major SV classes had the TRP/INV structures shown in Fig. 1 (top). Nine TRP/INVs were present, five of which mapped in the ∼1 Mb of ATE DNA and four dispersed in the remaining ∼ 36.5 Mb of genomic DNA. When compared to all single-read TRP/INV candidates derived from other cultures (G5, G10, G15), each TRP/INV was found only at the indicated stage of growth. Hence, within the limits of detection, TRP/INVs did not become fixed in the population and appeared to arise only transiently during growth (Fig. 1, bottom).

**Fig. 1.**
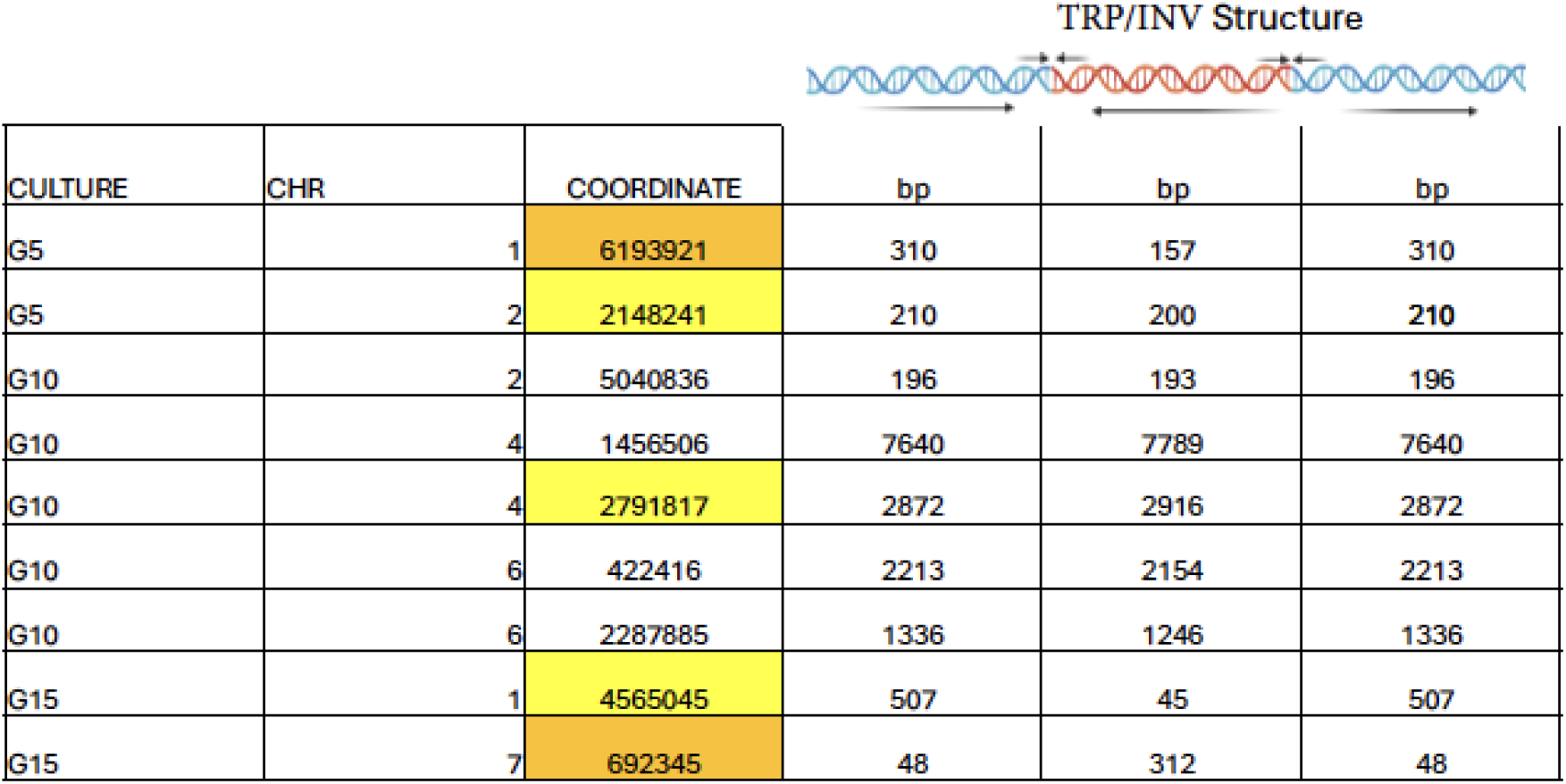
Initial Analysis of TRP/INV from Large Clustered Insertions in G5, G10 and G15 Cells. DNAs were isolated and sequenced after *A. flavus* NRRL3357 cells had been cultured for 5 (G5), 10 (G10), and 15 (G15) cycles, and subsequently screened for large (≥ 300 bp) insertions, using the Sniffles 2 algorithm as described in Materials and Methods with a variant read number of 3 and characterized further by IGV analysis. Individual reads were then analyzed by BLASTing to the G0 assembly to identify the TRP/INV shown here. The listed coordinates are the approximate sites of TRP/INV formation relative to the chromosomal (CHR) orientation. The yellow boxes refer to centromeric TRP/INV, while the orange boxes refer to subtelomeric TRP/INVs and TRP/INVs within ATE elements scattered at other sites in the genome. Uncolored boxes refer to TRP/INVs dispersed at non-ATE sites. The structure of the unprocessed (symmetric) TRP/INV is shown on top, with arrows below figure representing the relative orientations of TRP/INV direct and inverted repeats. Arrows above figure represent the typical palindromic sequences at these sites. The corresponding sizes of the direct (blue) and inverted (red) repeats are shown below.

### Quantitative Characterization of the TRP/INV Class in G15 Cultured Cells

To analyze the frequency of TRP/INVs quantitatively, we conducted an exhaustive study of all G15 SVs identified as ≥ 300 bp insertions (relative to the G0 assembly) that also were associated with a linked inversion. We characterized each of these reads and identified those having a TRP/INV configuration. These 221 TRP/INV species were distributed over all eight chromosomes of *A. flavus* (Table 2, SI_Table_S1). The low abundance and transitory nature of these species were further examined by quantifying all chromosome 1 TRP/INV in DNA derived from G0, G5, G10 and G15. We found three cases of heritable sites of TRP/INV. In each case, the TRP/INV structure was unique (SI_FigS1). We cannot rule out, however, the possibility that each of these had a common origin that subsequently underwent multiple rounds of rearrangement. These data reinforce the transient nature of TRP/INV.

**Table 2.** Chromosomal Distribution of G15 TRP/INV Among All Chromosomes.

| Chromosome | # TRP/INV |
| --- | --- |
| 1 | 33 |
| 2 | 23 |
| 3 | 44 |
| 4 | 27 |
| 5 | 34 |
| 6 | 14 |
| 7 | 20 |
| 8 | 26 |
| total | 221 |

Direct BLAST search of individual sequencing reads relative to the G0 telomere-to-telomere assembly revealed three TRP/INV subtypes (Fig. 2). Eighty-nine species fall into the symmetric class (class 1); Fig. 2, top) and have intact head-to-tail configurations at both inversion junctions within the estimated margin of error of sequencing and sequence alignment (Δ0-50 bp). One hundred thirteen species fall into an asymmetric class (class 2; Fig. 2, middle), displaying an intact head-to-tail configuration (Δ0-50 bp) at one of the two junctions and a more extensive deletion at the other end. In comparison, 19 species belong to another asymmetric class (class 3; Fig. 2, bottom) consisting of TRP/INVs containing a > 50 bp loss of sequence at both inversion junctions. SI_Fig_S2 shows examples of each of these classes. We also identified a series of complex forms that may be derived from multiple rounds of TRP/INV formation (data not shown).

**Fig. 2.**
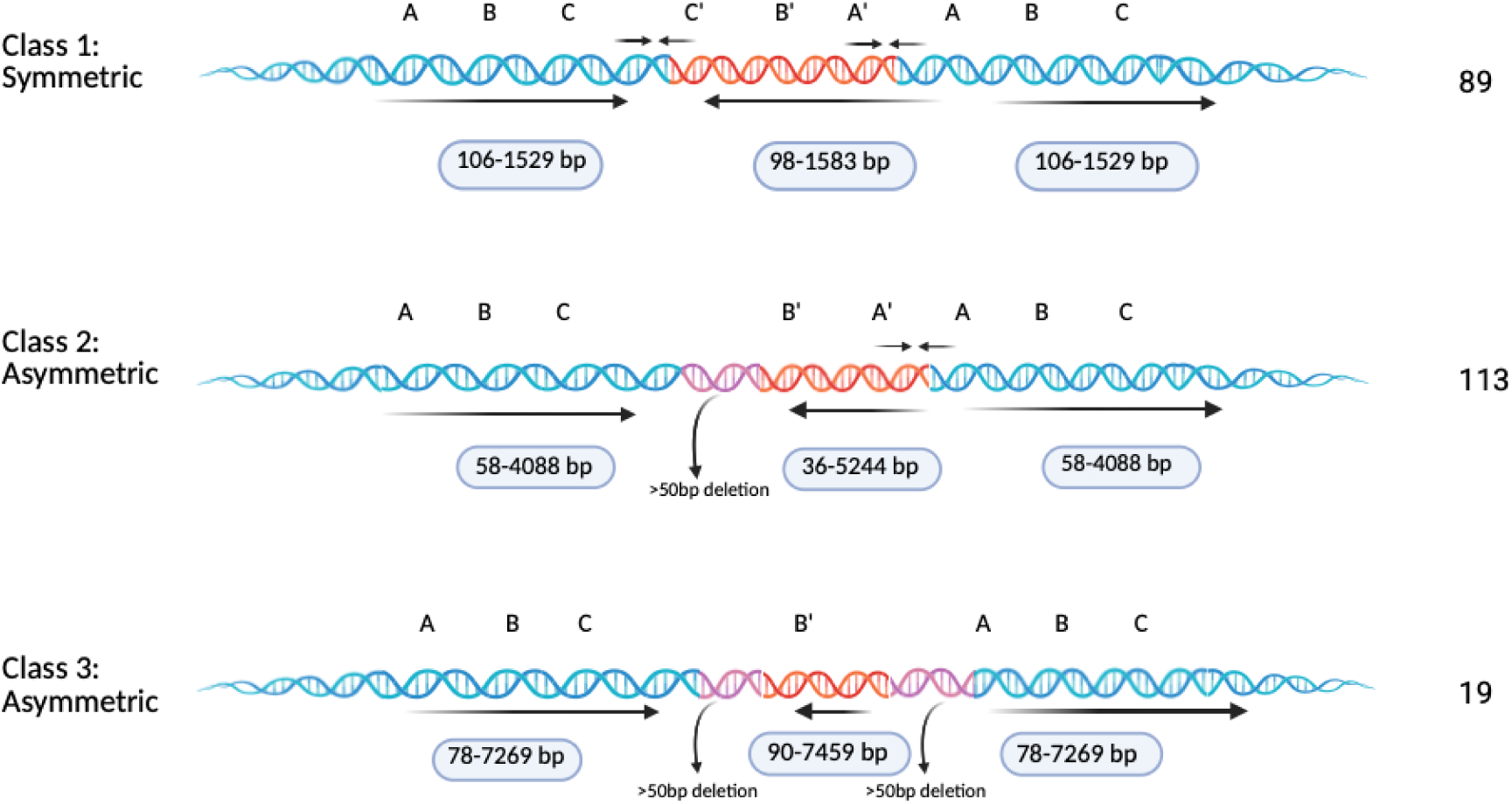
Three Structural Classes of TRP/INVs. The three classes of TRP/INVs deduced by BLAST analyses of nanopore sequencing queries to genomic assemblies are presented here. The first class (Class 1, symmetric) has little or no deletion at the junction between direct (blue) and inverted (red) repeats within the error of sequencing, set at ≤ 50 bp. Class 2 (asymmetric) has a deletion at one of the two junctions >50 bp. Class 3 (asymmetric) has deletions of >50 bp at both junctions. The size ranges of both direct and inverted repeats are listed below for each class and the number of observed species in each class is listed on the right. The orientation of hypothetical sequences A, B, and C (and complementary sequence A’, B’ and C’) as well as the directional arrows are shown for visual clarity as described in the legend to Fig. 1. Created in BioRender. Cary J. et al. (2025) https://BioRender.com/s70s676.

### TRP/INV Clusters at Centromeric and Subtelomeric ATEs

We analyzed the TRP/INV distribution to determine their potential enrichment in specific chromosomal domains. We found a 17.5-22-fold higher frequency of TRP/INV in the AT-rich elements (ATEs) present at centromeres (1/14.3 kb), subtelomeres (1/11 kb), and other classes of ATEs (1/11.5 kb), compared to the overall frequency at other genomic sites (1/251.6 kb) (Table 3A). The latter class distributes to both coding (101/145) and non-coding (44/145) regions (Fig. S2A-S2D; Table S1).

**Table 3.** Enrichment and Characterization of TRP/INVs in Centromeric and Subtelomeric ATEs. A) The chromosomal location, number, and fraction of TRP/INV in ATEs (centromeric, subtelomeric, and ATEs located elsewhere in the genome) and non-ATE (dispersed) regions, scattered among all chromosomes, are listed [24]. B) The direct and inverted repeat sizes (together with standard deviations) of TRP/INVs, located in centromeric and non-centromeric ATEs (subtelomeres + other ATEs) or dispersed non-ATEs are shown.

| Location | # TRP/INV | Frequency |
| --- | --- | --- |
| Centromeric | 56 | 1/14,285 bp |
| Subtelomeric ATE | 14 | 1/11,023 bp |
| “Other” ATE | 6 | 1/11,500 bp |
| Dispersed | 145 | 1/251,563 bp |

Table 3. **Enrichment and Characterization of TRP/INVs in Centromeric and Subtelomeric ATEs.**
| Location | Direct Repeat Sequence (bp) | Inverted Repeat Sequence (bp) |
| --- | --- | --- |
| Centromeres | 425 ± 202 | 400 ± 242 |
| Subtelomeres+ “Other” ATEs | 455 ± 367 | 421 ± 349 |
| Dispersed | 471 ± 733 | 498 ± 771 |

Analysis of the size of the direct and inverted TRP/INV sequences (Table 3B) revealed that the range of values was reduced (as shown by the standard deviation) in centromeres and subtelomeric (and other) ATEs, relative to dispersed non-ATE sequences. This reflects a tendency towards longer lengths in both the direct and inverted duplications in non-ATE TRP/INVs. For example, the lengths of such duplications extend up to 7.4 kb, compared to the maximum size of 0.92 kb at centromeres. Both the direct and inverted duplications in TRP/INV also trended to slightly smaller sizes in centromeric, compared to non-ATE, domains (Table 3B). These data suggest that selective forces may well restrict centromeric TRP/INV lengths.

We compared the TRP/INV patterns obtained from the analysis of the nanopore sequencing with the results from two independently PacBio (Revio) [34] sequenced G15 DNAs to ascertain the influence of sequence methodologies on the TRP/INV generation (SI_Table_S2). We observed qualitatively similar results with both sequencing methods, although TRP/INV levels were roughly 30-fold higher in ONT than in PacBio sequencing. Seven TRP/INVs were present, all at unique centromeric and subtelomeric sites. While substantiating the formation of TRP/INVs and their biased distribution towards AT-rich regions, these results suggest the presence of uncharacterized methodological biases in the library formation or sequencing of TRP/INV-containing DNA.

### Mitochondrial Genome Accumulates TRP/INVs Less Frequently Than the Nuclear Genome

As a control for the establishment of transient TRP/INVs and to test the contribution of library or sequencing anomalies that may generate these structures, we also measured SVs by the same methods in the high copy AT-rich mitochondrial DNA present in the in the sequencing reactions of each culture. The 30 kb circularly permuted and high copy mitochondrial genome is present in a sequencing depth of 20,000 to 30,000 copies. In contrast, centromeric sequences are in general covered in less depth (∼250 copies) than other parts of the genome. The mitochondrial genome therefore can serve as a model for the accumulation of TRP/INVs in centromeric sequences. In the G15 nanopore sequencing reads, we found only two TRP/INVs at unique sites. Due to the small sample size, we also explored the frequency of mitochondrial TRP/INVs in G0, G5 and G10 nanopore sequencing experiments. G0, G5, and G10 reads gave rise to 1, 2, and 0 TRP/INVs, respectively. If mitochondrial and nuclear TRP/INVs were found at a comparable frequency, we would have expected 210 TRP/INVs per 30 kb genome. Furthermore, the frequency of TRP/INVs in dispersed sequences is > 25-fold higher than in the mitochondrial genome, despite their similar palindrome density (Table 4). These results are inconsistent with a technical reason for TRP/INV formation and additionally suggest that aspects of nuclear DNA replication and recombination contribute to the formation of these species.

**Table 4.** Observed Frequencies of TRP/INVs at Centromeres and Mitochondrial Sequences. Out of the total of G0, G5, G10 and G15 DNAs (120 kb total) genomes sequenced, we found only TRP/INVs despite the higher depth of mitochondrial sequencing. Taking the predicted palindromes density of the 78% AT-rich mitochondrial genome into account, we estimate at least a 30-fold enrichment of TRP/INVs in the ∼800 kb of centromeric sequence and a >25-fold enrichment in dispersed nuclear sequences. Mitochondrial and centromeric palindromes were predicted using similar assumptions, as described in **Materials and Methods**. The origin of the sequenced genome is given in the parentheses.

|  | TRP/INV<br>(Observed) | Total Genome<br>Sequenced (kb) | Sequencing Depth<br>(Average) | TRP/INV Frequency<br>(/kb/read) | Predicted Palindrome<br>Density (/100kb) |
| --- | --- | --- | --- | --- | --- |
| **Centromere** | 56 | 800 (G15) | 278 | $2.5 \times 10^{-4}$ | 831 |
| **Mitochondria** | 5 | 120 (G0-G15) | 26140 | $1.6 \times 10^{-6}$ | 140 |
| **Dispersed** | 145 | 36500 (G15) | 1000 | $4.0 \times 10^{-5}$ | 100 |

### Identification of Palindrome Sequences at the Inverted/Direct Repeat Junction

Previous studies revealed that TRP/INVs of bacterial and yeast genes are formed via palindrome formation and template switching during bi-directional DNA replication or DNA repair which recreates the palindromic sequence in the symmetric product [8–10]. We tested the prediction that palindromes are at the site of TRP/INV formation, by identifying junctional read sequences that were complimentary to both strands of the genomic assembly by BLAST analyses (examples are shown in SI_Fig_S3A-S3D). The palindromes were then screened further for their predicted thermodynamic stabilities using the UNAfold mfold 4.2 program [35–37] (Table 5A-5C, Table 6). Seventy five junctions had palindromic characteristics and separate into those falling in centromeric (Table 5A), dispersed (Table 5B), and in subtelomeric (and other ATE) (Table 5C) locations.

**Table 5A.**
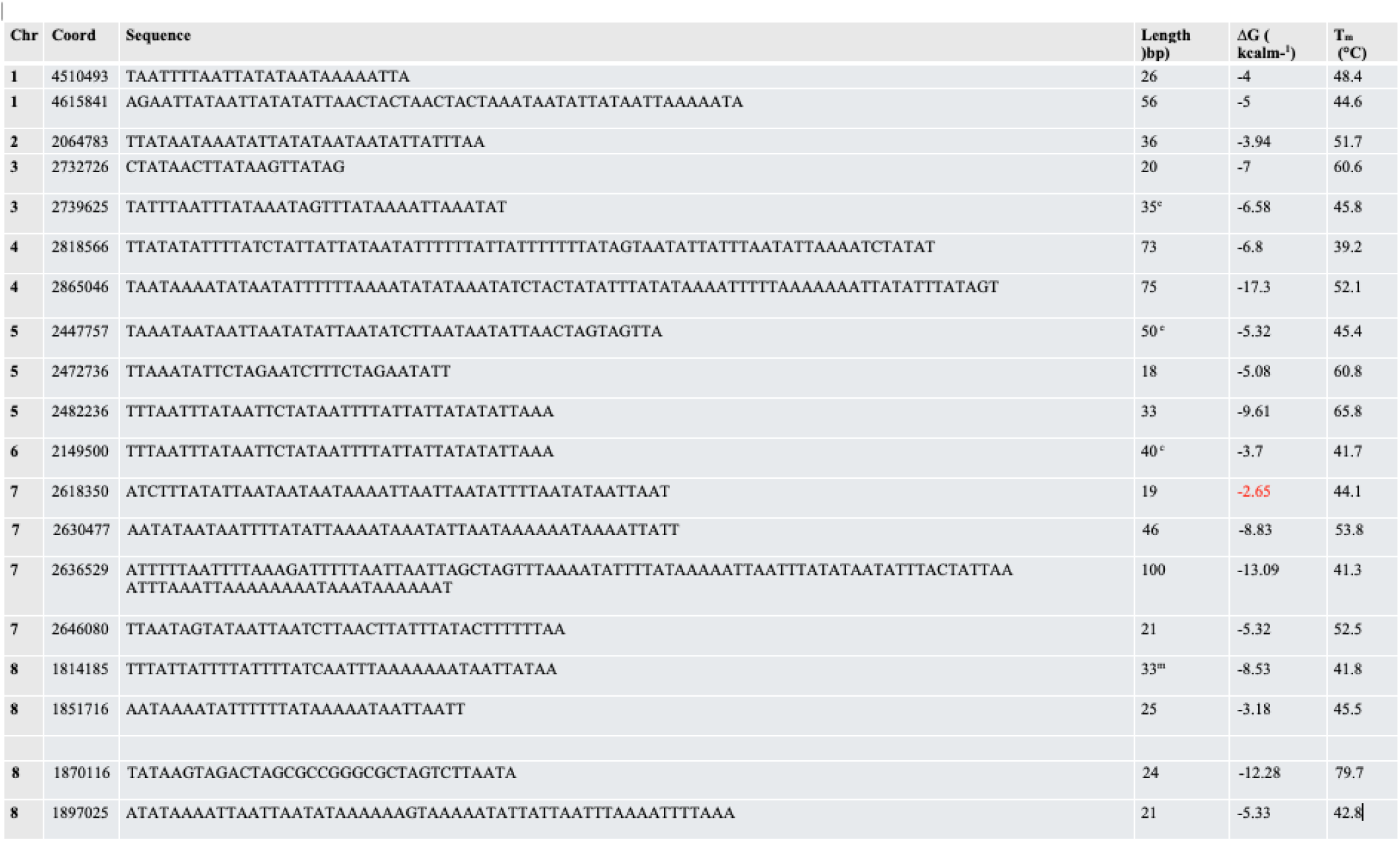
Predicted Junctional Palindromes in Centromeric TRP/INVs. The chromosomal coordinates, sequence coordinates, sequence, length and predicted ΔG (kcal/mol) and T_m_ values (°C) (calculated by UNAfold mfold 4.2 using the default settings: 25°C, 3 mM Mg, 50mM NaCl) of centromeric TRP/INV palindromes. Values of ΔG > -3 kcal/mole presented in red.

**Table 5B.**
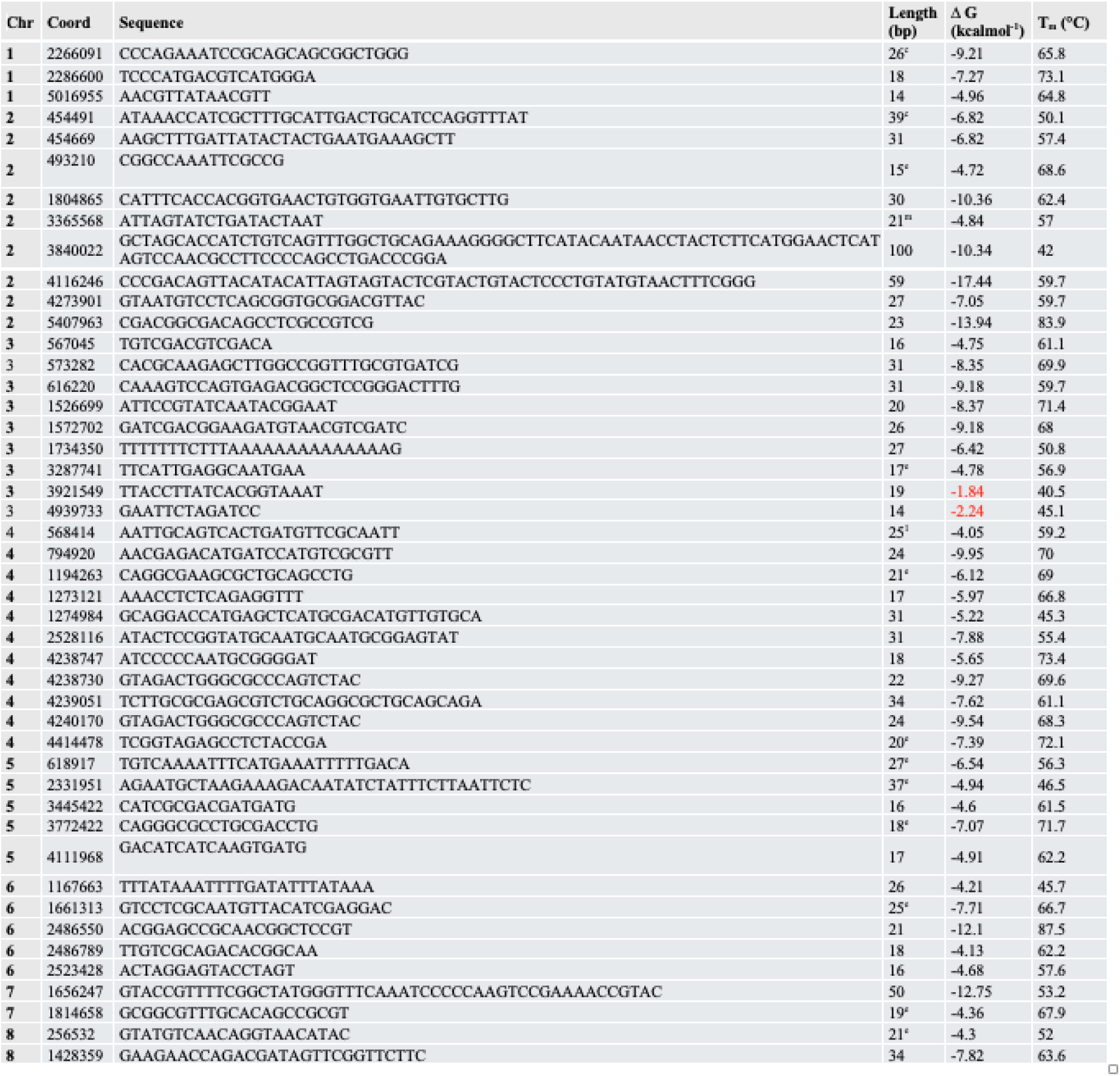
Predicted Palindromes at Dispersed Non-ATE TRP/INV Junctions. The chromosomal coordinates, sequences, length and predicted ΔG and T_m_ values of palindromes at inverted repeat junctions of non-ATE TRP/INVs. Stabilities were determined as in Table 5A. Those marked in red are > - 3 kcal/mole, calculated as in Table 5A. The superscript e refers to palindromes associated with extended regions of pairing; mp refers to multi-palindromic structures, and c refers to more complex pairing structures.

**Table 5C.**
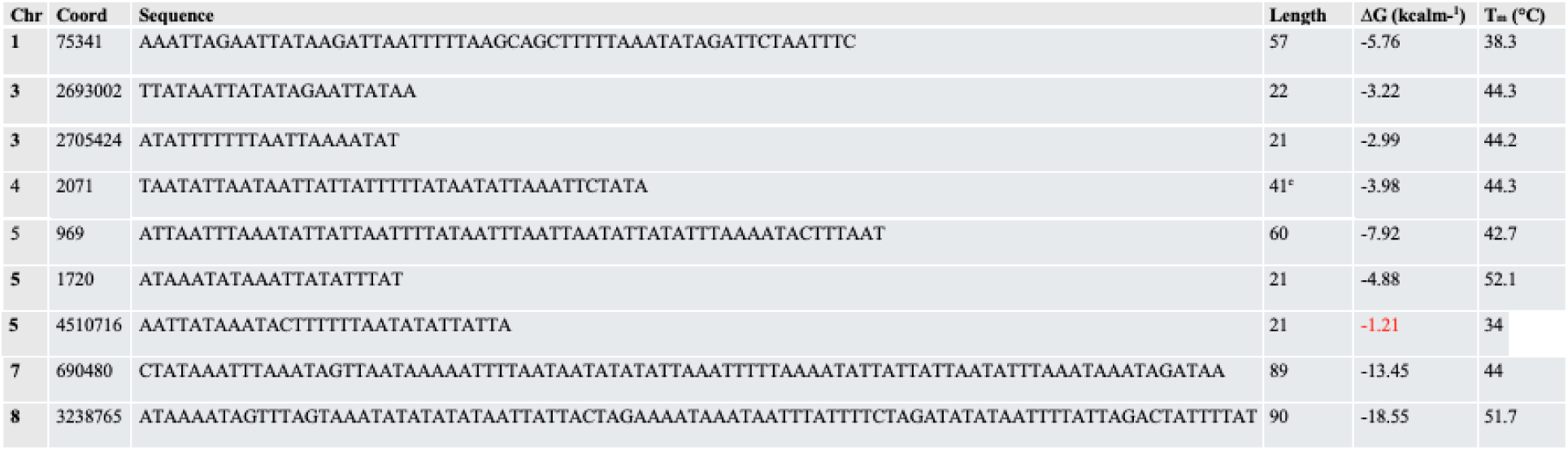
Predicted Palindromes at \Subtelomeric and Dispersed ATE TRP/INV Junctions. The chromosomal coordinates, sequences, length and predicted ΔG and T_m_ values of palindromes at inverted repeat junctions of subtelomeric (and other ATE) TRP/INVs of all stabilities were determined as in Fig. 5A. Those marked in red are > -3 kcal/mole (as calculated in Fig. 5A).

**Table 6.**
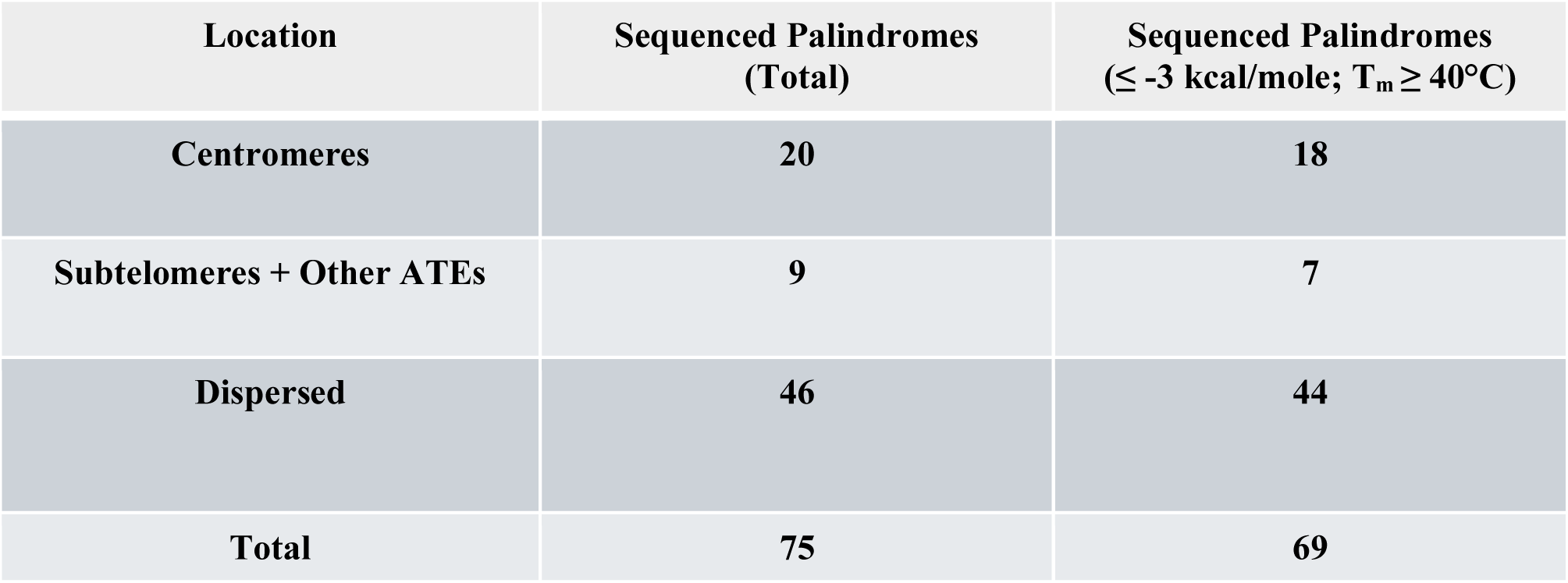
Palindrome Distribution Identified by BLAST and UNAfold Analyses. The distribution of palindromes from ATEs (centromeric and subtelomeric + ATEs located elsewhere in the genome) and non-ATE (Dispersed) regions are listed together with the distribution of palindromes predicted to have ΔG values of ≤ -3kcal and T_m_ ≥ 40°C. The palindromes of lower stabilities may still form secondary structures but do not meet the set limitations set in this study.

Palindromes ranged in size from 13-100 bp in non-ATE sequences and 18-100 bp in centromeric, subtelomeric (and other ATE) sequences. Based on the UNAfold DNA secondary structure algorithms, most putative palindromes have stabilities of -3 kcal/mmol or less (ranging from -3 to -20 kcal/mmol) and T_m_ values of at least 40°C (ranging from 40-84° C) (Table 5, right column). Based in part on the discussion below, -3 kcal/mol was used as the lowest stability cutoff if T_m_ were at or close to 40°C.

As a control, we compared the predicted stability of a previously isolated set of short palindromic junctions deduced to be at the sites of inverted sequence formation in yeast [7] (Table 7). These palindromes have ΔG values ranging from -2.31 to -7.06 kcal/mole and T_m_ values from 42-67°C, substantiating the validity of the choice of parameters in this study.

**Table 7.**
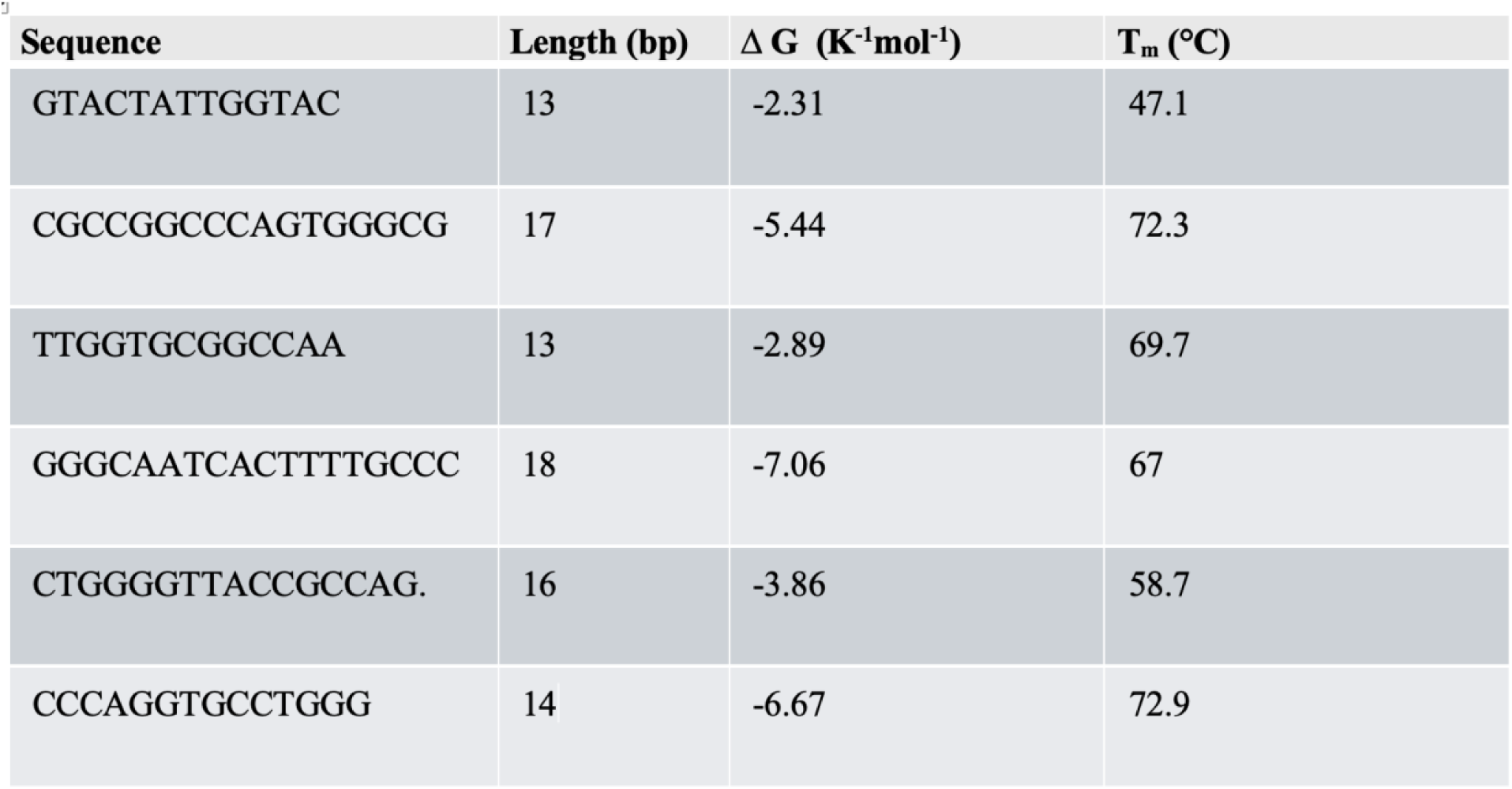
Predicted Stability of Palindromes Identified at Junctions of Inverted Repeats by Rattray et al (2005) [7].

Not all junctional sequences contained identifiable palindromic sequences within the threshold defined in this study. Out of 75 sequenced junctions with no significant deletion, four fell below the threshold value. This could either be an indication that the threshold is somewhat arbitrary or that a stable palindrome may not always be necessary for TRP/INV formation.

### Palindromes Are Present in Genomic Sequences at Sites Adjacent to the TRP/INV Junctions

We sought to test more generally whether palindromes are present in the genomic sequences that predict the formation of TRP/INVs. To this end, we examined the 200 bp genomic sequence surrounding the junction in unrearranged sequences at TRP/INVs sites that had junctional deletions of 40 bp or less, for the presence of simple palindromes, complex paired structures and multi-palindromic sequences. TRP/INV sites that contain larger deletions can obscure the actual site of palindromic sequences and were excluded from this study. We found that most junctions contain simple palindromes with or without extended associated pairing (147 [88%] in non-centromeric dispersed regions] 52 in centromeric regions [93%]; and 24 [100%] in subtelomeric domains). The palindrome lengths (stem plus loop) averaged 33 bp in dispersed regions, and 41-43 bp in AT-rich subtelomeric and centromeric regions. All stabilities averaged -6.7 kcal/mol, while T_m_ values averaged higher in dispersed (56.4°C) than in AT rich regions (48°C), as expected due to differences in base composition [Fig. 3; (SI Table 3A-C)]. A fraction (10%) of junctions had stabilities with ΔG > 3.0 kcal/mol. Some junctions were associated with long regions of extensive pairing (13%), and a fraction (12%) had more complex pairing patterns or multi-palindromic structures (SI Fig. 4 for examples). These findings raise the possibility that some homologous pairing patterns may have a role in perturbing localized structure with or without strand exchange. These predicted structures are unlikely to be static. Rather, they are likely to represent one among many in an equilibrium of multiple palindromic structures.

**Fig. 3.**
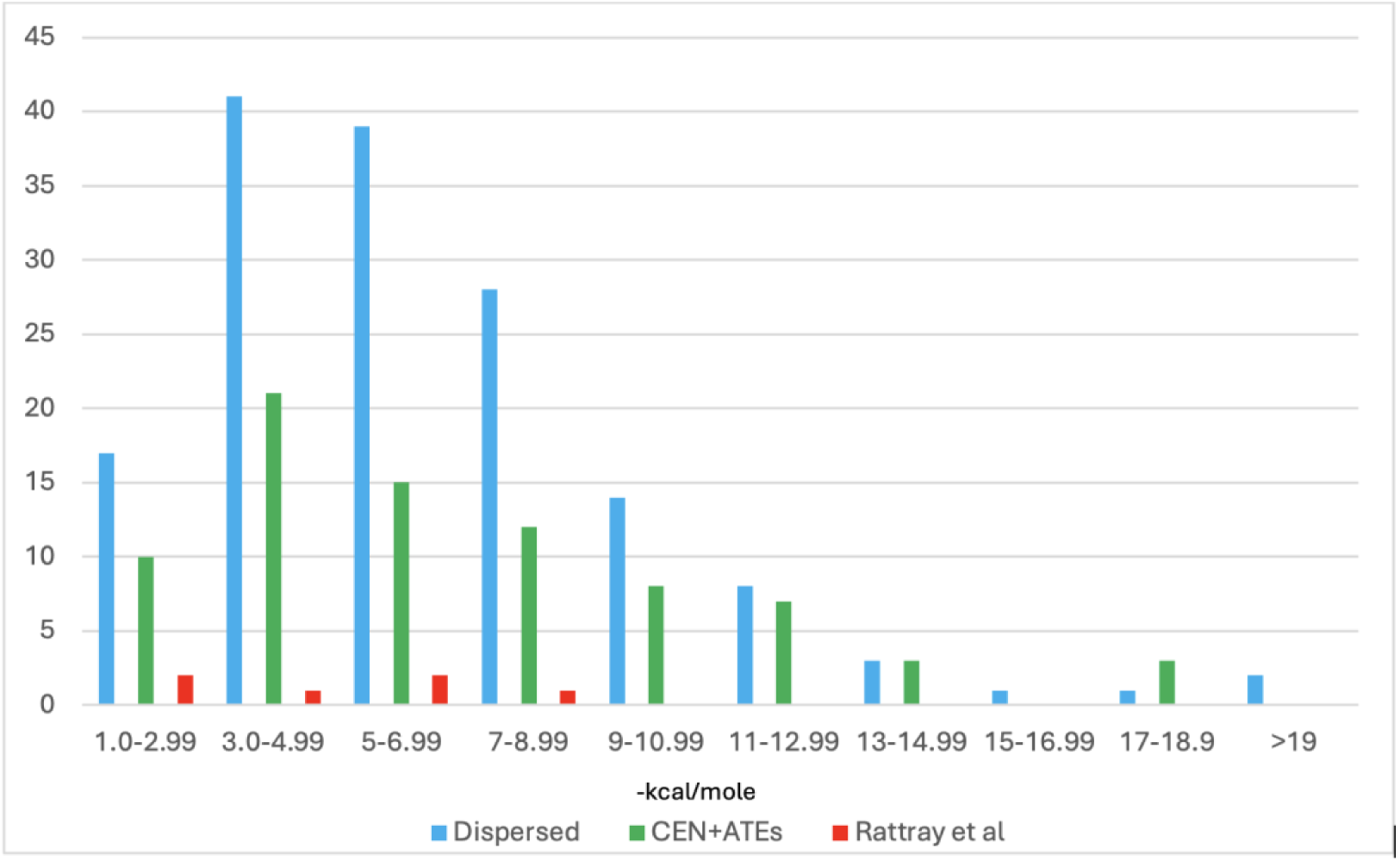
Palindrome Stabilities in Centromeric, Dispersed ATE and Non-ATE Sequences. Palindrome Stabilities Predicted by UNAfold Analyses in TRP/INVs with a deletion of 40 bp or less. The distribution of TRP/INV palindromes in centromeric, subtelomeric, and other ATEs (CEN + ATEs) (green) and from dispersed non-centromeric (Dispersed) sequences (blue) are shown as a function of stability (-kcal/mol) of as determined by UNAfold analysis. Also shown are the stabilities found at sites of inversion in yeast (Rattray et al MCB 2006) (red) for comparison.

**Fig. 4.**
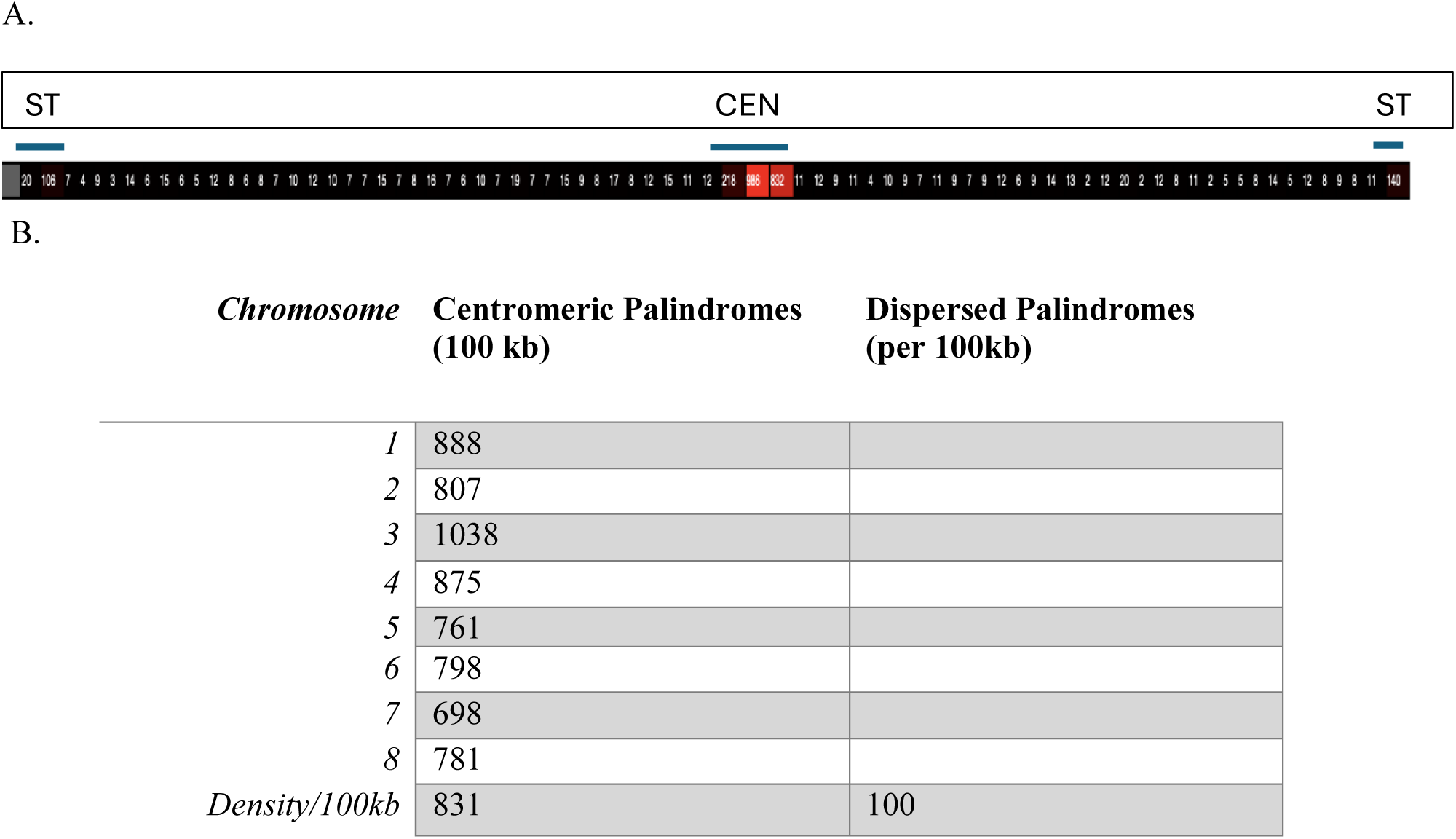
Predicted Distribution of TRP/INV-Associated Palindromes. (A) Heat Map of Predicted Distribution of Centromeric TRP/INV-Associated Palindromes (before ΔG filtering) (B) Predicted Centromeric TRP/INV-Associated Centromeric Palindromes are Present at Higher Densities than Dispersed TRP/INV Associated Palindromes (measured on a segment of ca 60 kb segment of chromosome 3). The assumptions used for calculations of these values are described in the text. CEN, centromere; ST, subtelomeric.

**Fig. 5.**
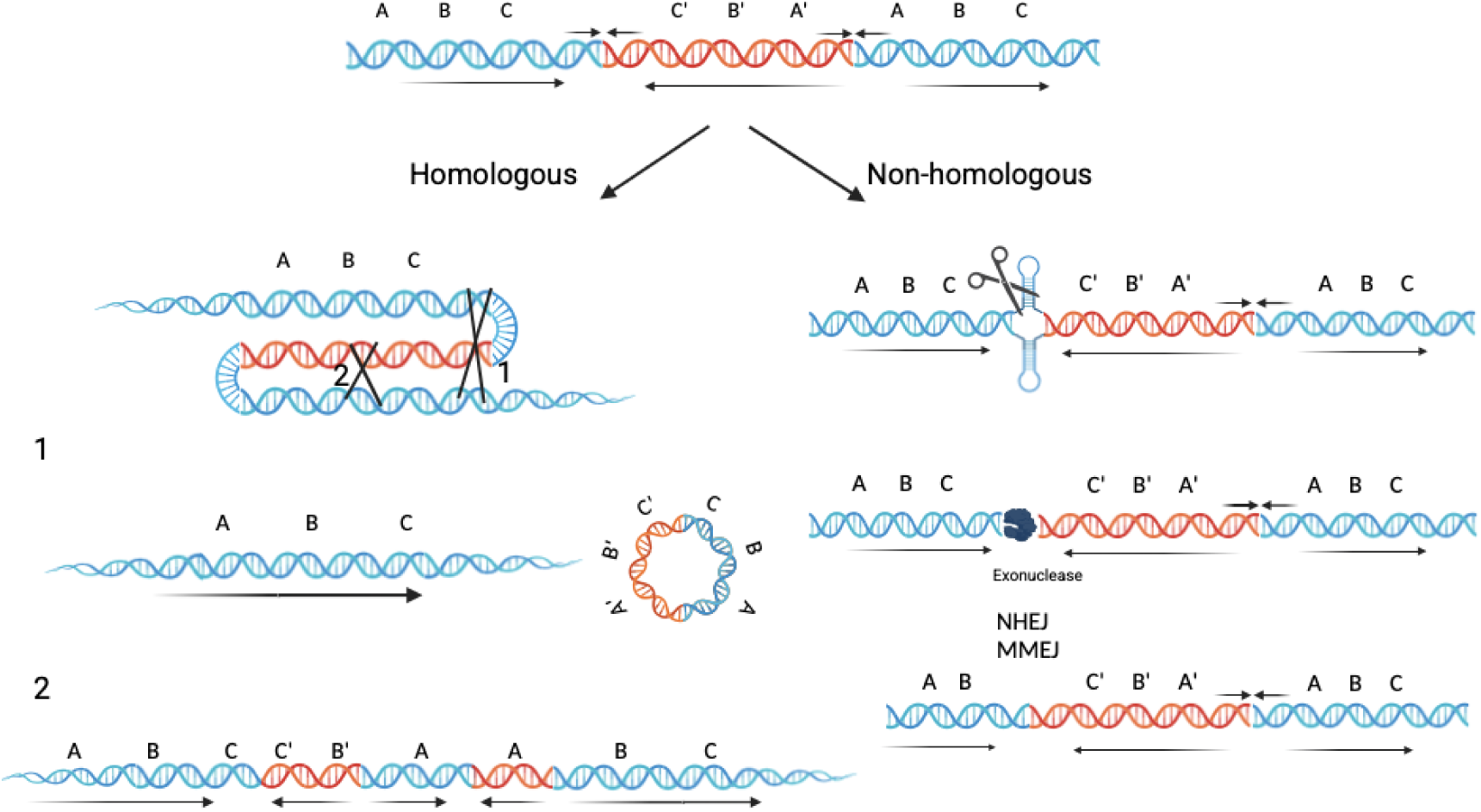
Models for the Resolution of TRP/INV Structures. Homologous and non-homologous pathways for resolution of TRP/INVs. (Top) The structure of a TRP/INV. Color designations are described in Fig. 1. (Left) Recombination between direct repeats (1) gives rise to excision of a dimeric inverted circle that has the potential to reintegrate into the genome, creating repeat amplifications, while recombination between direct and inverted repeats (2) creates additional inversions that undergo further rearrangements. (Right) The extrusion of a palindrome is expected to undergo resolution or cleavage, resulting in a double strand break susceptible to exonuclease digestion. Microhomology mediated end joining (MMEJ) or non-homology end joining (NHEJ) can then produce Class 2 and Class 3 TRP/INV deletions. The orientation of hypothetical sequences A, B, and C and the directional arrows are shown for visual clarity. Created in BioRender. Cary, J.W. et al (2025) https://BioRender.com/r14e12.8

### Palindrome Formation Mirrors TRP/INV Centromere Enrichment

Regions of high AT or GC content produce a greater number of palindromes on a simple statistical basis, and we observed an even greater frequency of palindromes in centromeric, subtelomeric and other ATEs (Table 1). We wanted to test whether the frequency of a palindrome, having the predicted characteristics of TRP/INV-associated palindromes, could be sufficient to explain the higher density of TRP/INVs at centromeres. We therefore estimated the relative number of palindromes that would be likely to generate a TRP/INV in centromeric and dispersed regions (see **Materials and Methods**). To this end, we calculated the most frequently observed stem and loop sizes, as well as ΔG and ΔG_cf_-ΔG_lin_ values, the latter a measure of the energetic favorability of cruciform formation [38]. Due to the complexity of the centromeric palindrome patterns, we first determined that the most frequent classes of the terminal portions of the TRP/INV-associated palindromes in the centromere of chromosome 3 fell into a stem size of 12-100 bp, a loop size of 4-8 bp with 0-2 stem mismatches. The values were input into a program, Palindrome Analyzer [39], designed to estimate the frequency and stabilities of palindromes at the chromosomal level, and the sites of the TRP/INVs were confirmed to be present in the output. The distribution of utilized palindromes had ΔG_cf_-ΔG_lin_ values that fell between 10 and 16 kcal/mol, which were applied proportionally to the output, since the distribution of stabilities in the observed and predicted palindromes were quite similar. The frequency of the predicted palindromes was then corrected to account the number of false positives. Based on this analysis, we estimate an average 100 kb region of centromere contained on average over 800 palindromes (Fig. 4). However, these values could not be applied to non-centromeric sequence due to the differing TRP/INV-associated palindrome parameters in AT-rich and dispersed regions. Examination of non-centromeric TRP/INV-associated palindromes in chromosomes 1, 3 and 4 revealed stem lengths of 5-12 bp, loop sizes of 4-19 bp, and stem mismatches of 0-1. In this case, the observed and predicted patterns were significantly different, with ΔG_cf_-ΔG_lin_ values of the observed palindromes distributed as follows: 15 at 3.1-6.0 kcal/mol (Class 1), 10 at 6.1-9.0 kcal/mol (Class 2) and 14 from at - 9.1-12.0 kcal/mol (Class 3). Since, the predicted TRP/INV-associated palindrome output had a different relative distribution within a 60 kb region of centromere 3 (24 in class 1, 105 in class 2, and 507 in class 3), we normalized the observed recombination frequencies using a weighted eligibility model [40, 41] (**see Materials and Methods**). Under these conditions, Palindrome Analyzer predicted ∼100 palindromes/100 kb. Although these values reflect the characteristics of most, but not all, palindromes, the overall frequency of palindromes clearly mirrors the enrichment of TRP/INVs in centromeric regions. We propose therefore that the abundance of short palindromes is a major driver of TRP/INV formation.

## DISCUSSION

Here, using the *A. flavus* model system, we report the first evidence, to our knowledge, for spontaneously formed TRP/INVs, a process is driven by the frequency of palindrome formation. Several lines of evidence support the identification of spontaneous but infrequent inverted triplications in *Aspergillus flavus* through structural variant analysis following high-coverage third-generation sequencing. Although we did not apply any intentional selection during growth, selective pressures induced by vegetative growth are possible. Indeed, in studies to be presented elsewhere, we found reductions in conidiation levels between the fifth and tenth weeks of culturing (data not shown).

First, BLAST searches of individual reads of presumptive large insertions generated after two independent methods of genome sequencing (ONT and PacBio) predicted TRP/INV structures consistent with unprocessed symmetric and processed asymmetric TRP/INV forms [8–10], albeit differing in abundance. Asymmetric products with deletions at one or more junctions are consistent with a role for a classical or alternative non-homologous end joining (see below). A large fraction of sequenced inversion junctions and sequences in short, deleted regions of asymmetric junctions revealed palindromes and quasi-palindromes at the site of genomic rearrangement, arguing for a mechanism that favors the use of a hairpin structure to generate the inverted sequence.

Second, we have shown that TRP/INVs are clustered at centromeres and centromeric AT-rich DNA. Interestingly, the abundance of palindromes in centromeric regions mirror the clustering of TRP/INVs. In addition, these studies have found that palindrome formation (as well as direct and inverted repeats) is also significantly elevated in genomic, compared to randomly generated DNA having the same AT content. These data argue for a central role of palindromes in driving TRP/INV formation and clustering in centromeric and subtelomeric regions. However, palindromes and AT-rich DNA are likely insufficient for TRP/INVs since the mitochondrial genome has a significantly decreased TRP/INV frequency per unit size, despite being AT-rich. In addition, mitochondrial TRP/INVs were present at a lower abundance than in dispersed sequences despite having a comparable density of predicted TRP/INV-associated palindromes. These data argue for a preference for inverted triplications in the nuclear genome.

Third, we found that some palindromes were predicted to be associated with more extended regions of homology or had more complex pairing structures. Although it is likely that predicted algorithms only reflect a snapshot of likely stable forms, the data does raise the interesting possibility that palindromes may have functions beyond or in addition to strand transfer [required for template switching (see below)], such as in structural perturbations contributing to fork stalling.

Fourth, we found that the TRP/INVs are infrequent and do not become fixed in the population after 15 cycles of forced vegetative growth. For the G15 cultures sequenced by ONT, we identified 221 reads out of 3.6 x 10^6^ reads of sizes (> 2kb) that were used in this analysis. Since each read is derived from an independent DNA molecule, this suggests an overall frequency of 6 x 10^-5^ TRP/INV/genome. This estimate is only an approximation, since the coverage of different genomic regions is not identical using either sequence method. These structures arise at every stage of subculturing and are in low abundance, possibly due to high rates of TRP/INV processing.

Such TRP/INV instability may arise via distinct mechanisms of homologous and non-homologous recombination (Fig. 3 left). Recombination between the direct repeats would result in excision of a TRP/INV extrachromosomal circle, restoring the original structure (Fig. 3, left, pathway 1). On the other hand, recombination between inverted repeats would generate a novel pattern of direct repeats and inversions (Fig.3, left, pathway 2). Indeed, the high frequency of direct and inverted repeats that we have found in centromeric and subtelomeric regions are among the expected products of TRP/INV recombination. In contrast, unequal recombination between sister chromatids would result in both the loss and duplication of these regions.

The instability of palindrome DNA that leads to cruciform and hairpin extrusion and promotes double strand break formation has been well-documented in both prokaryotes and eukaryotes [4]. Several pathways may be responsible for the resolution of the double-strand break, including exonucleolytic degradation coupled with microhomology-dependent non-homologous end-joining (MMEJ) [42, 43] (Fig. 3 right), or alternatively, Ku-dependent non-homologous end-joining (NHEJ) [43]. MMEJ is indeed found in yeast and filamentous fungi, although by mechanisms independent from the pol θ-dependent process in multicellular organisms and is likely to involve, in part, subunit 3 *(S. pombe cdc27*, S*. cerevisiae pol32*) of polymerase ∂ [42, 44]. The frequent processing of palindromic DNA in AT-rich sequences coupled with MMEJ or NHEJ is also expected to generate small deletions within these regions. This means of resolution is likely to account for characteristics of asymmetric class 2 and 3 TRP/INVs.

The transient presence of TRP/INV could also be due to a strong selection against the rearranged sequences leading to diminished growth rate or cell death. This explanation seems unlikely, however, given the multinucleate nature of mycelial growth in *A. flavus*, which would permit the retention of deleterious rearrangements [28]. If such a selection is present, however, it is more likely to be mediated through TRP/INV disruption of coding regions.

While TRP/INV-like structures were identified in the centromeric and subtelomeric regions using nanopore and PacBio sequencing, the frequencies of TRP/INV formation differ between the two methods [30, 34]. Quantitative analysis using such different methodologies is indeed problematic. Although TRP/INVs are unlikely to be the consequence of any known sequencing artifacts [45], as supported by the reduced frequence of TRP/INV formation in mitochondrial DNA, several methodological distinctions could contribute to these quantitative differences. For example, nanopore sequencing results in a direct readout of the structural variant, while PacBio amplifies each extracted read to provide greater consistency and potentially selects against TRP/INV amplification. In addition, the pre-treatment methods to repair damaged DNA before the construction of the appropriate library may bias the size of DNA fragments in distinct ways. We cannot, unfortunately, test these speculative possibilities at present.

Our data suggest that both structural and strand transfer properties of palindromes may be involved in the formation of TRP/INV structures. The mechanism of formation is likely to be akin to the models from Roth’s laboratory [8] for TRP/INVs in *Salmonella* (Fig. 6). We propose that the structural constraints of palindromes result in fork stalling, followed by subsequent palindrome resolution forming a foldback that primes the replication of the inverted repeat. Processing of the second palindrome is then responsible for the synthesis of the direct repeat. Replication stalling has indeed been observed in experimentally manipulated constructs in both prokaryotes and eukaryotes [12, 13]. Following palindrome-mediated template switching, microhomology-mediated pathway, such MMEJ or “micro” single-strand annealing (SSA) processes, would prime the synthesis of the direct repeat [46]. This model is appealing, since we can experimentally test the genetic dependence on proteins involved in homologous recombination, MMEJ and micro-SSA, as well as the inhibition of these processes by Ku binding and protection of damaged DNA. The presence of palindromes formed in the TRP/INV product can also give rise to multiple cycles of inversion and duplication, leading to the observed complex products (data not shown).

**Fig. 6.**
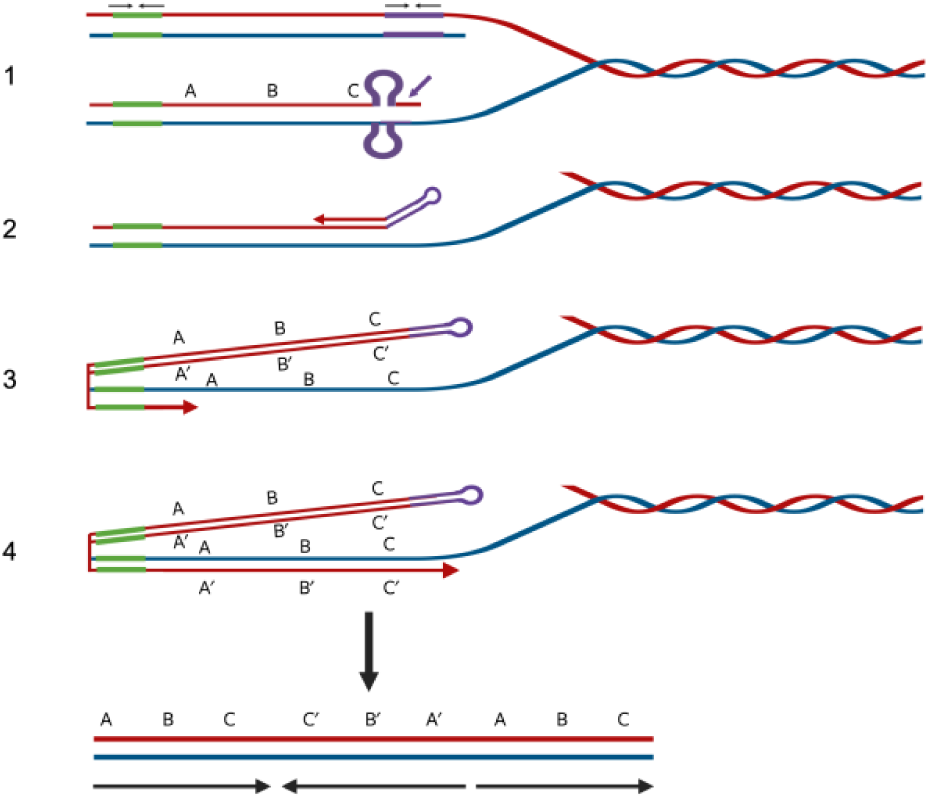
A Mechanism for Palindromic and Direct Repeat Strand Exchange in TRP/INV Formation. A four-step model modified from Roth and colleagues [8], based on their extensive studies in *Salmonella* is presented: 1. Palindrome extrusion and processing in replicating DNA and resolution by nucleolytic activities. 2. Formation of hairpin foldback that primes synthesis of the inverted repeat. 3. DNA synthesis reaches the second palindrome (green) followed by Ku-independent (or inhibited) mediated strand transfer and the priming of the direct repeat synthesis(4). Created in BioRender. Cary J.W. et al. (2025) https://BioRender.com/m61z558

One of the exciting aspects of this study is the accumulation of TRP/INV in AT-rich subtelomeric and centromeric sequences which is due, in part, to the critical role of palindromes abundance in TRP/INV formation. Indeed, this study is the first characterization of short repeat-mediated recombination within centromeric and subtelomeric regions. This finding may be related to our previous observation that AT-rich regions at the centromeres of non-homologs (and in many cases, even homologs) are highly divergent. We also found that specific subtelomeric ATE sequences are present at different sites within and among strains. This observation may be related to the role of subtelomeres in the elimination or the evolution of secondary metabolite gene clusters and their participation in other selected rearrangements in fungi [25]. Both results support an equilibrium model, whereby deleterious functions such as perturbation of kinetochore function or rampant subtelomeric rearrangement may counteract evolutionary advantageous roles for centromeric divergence and subtelomeric sequence shuffling. We propose that mechanisms that regulate such an equilibrium must be present at centromeres and subtelomeres, possibly modulated by recombination activities. In addition, heterochromatin formation may be regulated. The fungal centromeric (and probably subtelomeric) AT-rich sequences are heterochromatic. In yeast, as in most eukaryotes, histone modification and chromatin condensation lead to reduced levels of transcription, recombination and DNA repair in heterochromatin [47–49]. For example, the histone deacetylase protein Sir2 is responsible for telomeric heterochromatin condensation and position effects [50]. The use of *A. flavus sir2* (*sirE*) mutants with wide-ranging phenotypes [51] is an attractive possibility to test this hypothesis. Such regulation may be the underlying basis for the selection against larger sizes of centromeric TRP/INVs. In addition, subtelomeric genes that are bounded by heterochromatic regions may serve to regulate transcription from these regions.

Our findings suggest that centromeres and subtelomeres can be more recombinationally active than anticipated. ATEs are maintained in all filamentous fungi and are indeed sites of evolutionary polymorphism [20]. However, any region that is high in AT or GC content has a statistically higher probability of palindrome formation and may be subject to a similar process, a possibility that requires further exploration. The TRP/INVs reported here are yet another example of the reported association of AT-rich sequences with genetic instability [52–54], which may form hot spots for rearrangement in the eukaryotic genome in normal and diseased cells. The exploration of spontaneous TRP/INVs in other genetically manipulatable organisms using the methods described above, will be a valuable future direction to test the generality of this finding.

TRP/INVs are associated with genetic diseases. Brewer et al. (2024) conducted an extensive study that demonstrated that many human disease-related loci had structures consistent with the involvement of TRP/INVs, and these loci were often flanked by direct repeats [14]. Other laboratories have also demonstrated that these structures are one of the most prevalent mutational signatures in cancer and a common source of disease-related genomic instability [55, 56]. Understanding the fundamental mechanisms of TRP/INV formation in the absence of selection will be critical to understand the underlying basis of genetic instability that precedes the chromosomal amplifications found in these disease states.

## MATERIALS AND METHODS

### Strains, media, and growth conditions

G0 spore stocks of *Aspergillus flavus* NRRL 3357 were generated following nine-point inoculations of a diluted 1x10^3^/ml spore stock onto the surface of double-strength 5/2 agar [(per liter: 50 mL V8 juice, 40 g agar, pH 5.2 [57, 58]] and incubation at 30 °C in the light for five days, a condition that promotes conidiation. NRRL 3357 conidia were collected from plates by gentle scraping of the agar surface with 10 ml sterile 0.01% Triton X-100, then transferred to a sterile 15 ml screwcap polypropylene tube, and stored at 4°C. For forced vegetative growth studies, addition of 1 ml (about 1x10^8^) of the spore suspension into 100 ml of A&M medium (per liter: sucrose, 50 g; (NH_4_)_2_SO_4_, 3 g; KH_2_PO_4_, 10 g; MgSO_4_ ·7H_2_O, 2 g; Na_2_B_4_O_7_ · 10H_2_O, 0.7 mg; (NH_4_)_6_Mo_7_O_24_ · 4H_2_O, 0.5 mg; Fe_2_(SO_4_)_3_ · 6H_2_O, 10 mg; CuSO_4_ · 5H_2_O, 0.3 mg; MnSO_4_ · H_2_O, 0.11 mg; and ZnSO_4_ · 7H_2_0, 17.6 mg.; pH 6.5) and incubation at 28 °C, at 200 rpm, in the dark for seven days, led to generation 1 of the experiment (G1). After 7 days (d), 20 ml of the G1 culture was removed from the 100 ml A&M shake culture and placed in a sterile 50 ml screw cap, polypropylene tube, then macerated for 30 sec using the Tekmar Tissuemizer (Tekmar Co., Cincinnati, OH). One ml of macerated G1 mycelia was transferred into fresh 100 ml A&M broth in a sterile 250 ml Erlenmeyer flask and incubated for another 7 d. This culture represented the second “generation” (G2) of vegetative growth. This process was continued for 15 cycles (15 weeks; 105 d). We diluted a 10 µl aliquot of macerated mycelia in 990 µl sterile water, spread 10 and 100 µl aliquots onto the surface of double-strength 5/2 medium, and incubated for three days to observe colony phenotype. Mycelia for genomic DNA preparation from 5 (G5), 10 (G10), and 15-week (G15) cultures of NRRL 3357 mycelia were collected as follows. At the end of the desired growth period, 50 ml of potato dextrose broth (PDB, Difco) was inoculated with 0.5 ml of macerated mycelia and incubated for 24 h at 28 °C with agitation (200 rpm). For NRRL 3357 generation 0 (G0), genomic DNA was prepared from a 24 h 50 ml PDB culture that had been inoculated with 0.5 ml of a 1x10^8^ spores/ml NRRL 3357 stock. Mycelia were collected from the 50 ml PDB shake cultures by vacuum filtration through sterile miracloth (EMD Millipore, Temecula, CA). The remaining liquid was squeezed from the collected mycelial mat by pressing it between paper towels. Mycelia were lyophilized for at least 24 h and stored at -80 °C until needed.

### Preparation of genomic DNA

Genomic DNA was prepared from lyophilized mycelia using the Qiagen Plant Mini kit (Qiagen, Beverly, MA) following the manufacturer’s protocol. Lyophilized mycelia were ground to a fine powder in liquid nitrogen and stored at -80 °C until needed. Briefly, 15 mg of each ground sample was processed in duplicate. After a spin column, the DNA was eluted with two 100 µl elutions using Qiagen AE buffer. The DNA suspension was further enriched by the addition of an equal volume of 25:24:1 phenol: chloroform: isoamyl alcohol (IAA). The upper layer was collected and extracted with an equal volume of 24:1 chloroform: IAA, and then precipitated by the addition of one-tenth volume of 3M sodium acetate pH 5.2 solution and two volumes of cold 95% ethanol followed by centrifuging at 10K rpm for 10 min. The DNA pellet was washed with 70 % ethanol, dried in a speed vac, and resuspended in 50 µl TE. DNA quantity and quality were determined using a NanoDrop OneC (Thermo Scientific). The quality of genomic DNA was also determined by electrophoresis on a 0.7% agarose gel.

### Sequencing

CD Genomics (Shirley, NY) carried out all sequencing. Both sequencing methods gave an average of 500-1500-fold coverage, although not equally distributed over all parts of the genome.

#### Nanopore sequencing

The library was prepared using the SQK-LSK112 ligation kit under standard conditions. The purified library was loaded onto primed R10.4 Spot-On Flow Cells and sequenced using a PromethION sequencer (Oxford Nanopore Technologies, Oxford, UK) with 48-h runs [30]. Base-calling of raw data used the Oxford Nanopore Dorado v 0.7.3 (https://github.com/nanoporetech/dorado/releases).

#### PacBio HiFi Sequencing

The SMRTbell library was constructed using the SMRTbell Express Template Prep kit 2.0 (Pacific Biosciences). Briefly, ∼2μg of the genomic DNA was carried into the first enzymatic reaction to remove single-stranded overhangs followed by treatment with repair enzymes to repair any damage that may be present on the DNA backbone. After DNA damage repair, ends of the double-stranded fragments were polished and subsequently tailed with an A-overhang. Ligation with T-overhang SMRTbell adapters was performed at 20° C for 60 minutes. Following ligation, the SMRTbell library was purified with 1X AMPure PB beads. The size distribution and concentration of the library were assessed using the FEMTO Pulse automated pulsed-field capillary electrophoresis (Agilent Technologies, Wilmington, DE) and the Qubit 3.0 Fluorometer (Life Technologies, Carlsbad, CA). Following library characterization, 3μg was subjected to a size selection step using the BluePippin system (Sage Science, Beverly, MA) to remove SMRTbells ≤15 kb. After size selection, the library was purified with 1X AMPure PB beads. Library size and quantity were assessed using the FEMTO Pulse and the Qubit dsDNA HS reagents Assay kits. The sequencing primer and Sequel II DNA Polymerase were annealed and bound, respectively, to the final SMRTbell library. The library was sequenced using 8M SMRT Cell on the Sequel II System with Sequel II Sequencing Kit [34].

### Assemblies

We bootstrapped G0 and G15 assemblies using both *A. flavus* strain NRRL-3357 (GCA_009017415.1_ASM901741v1) [23] and mitochondrial (*A. flavus* SRRC1009) [59] sequences that were concatenated to prepare a genome fasta file. This file served as input for G0 ONT or G15 PacBio reads to Rebaler (https://github.com/rrwick/Rebaler) [60]. G0 and G15 assemblies, stored in the Sequence Server Cloud, are available upon request.

### Structural variant determination

Structural variants were identified relative to the G0 assembly using Sniffles 2 (version 2.2) [32]. Alignments used minimap 2 [33] (version 2.26-r1175) with a gap open and gap extension set at 16 and 2, respectively. This version was determined empirically to be only ∼70% efficient for low abundance sequences in simple sequence regions, leading to both false positives and negatives. We therefore often coupled the algorithm with visual confirmation using IGV (version 2. (10.0)) [61, 62].

For qualitative analyses, structural variants were identified as ≥ 300 bp insertions with variant read numbers (DV) of 3 and confirmed by IGV analysis. Individual reads from these nanopore sequenced DNA were then used as BLAST queries to the G15 or G0 assemblies using SequenceServer (London, UK) [62]. We analyzed simple sequence organization for AT-rich sequences by removing the dust filter.

For quantitative analysis, we selected all reads that had a DV of 1 and individual reads from these nanopore sequenced DNAs were then analyzed by BLAST as noted above. A similar analysis was carried out for PacBio insertions of ≥ 200 bp. For all studies, the variant frequency (VAF) calculated as (variant read number (DV)/reference read number (DR) x 100). VAF values ranged from 0.1 to 0.2 %,consistent with a high coverage.

The most frequent products identified by BLAST analysis among large “insertions” were duplications and tandem inverted duplications. The TRP/INV species were categorized based on the extent of deletion at each junction between duplicated and inverted sequences. We established 50 bp as the maximum error at a junction, likely due to sequencing and alignment defects.

### Palindrome analysis

Palindromes were identified first from BLAST analysis of single reads with the G15 assembly by the presence of sequences at inverted sequence junctions homologous to both strands of the genomic sequence, representing the palindromes that initiate TRP/INV formation. We determined the predicted relative thermodynamic stability (T_m_, ΔG, ΔH, and ΔS) and secondary structure predictions of the palindromes by the application of UNAfold mfold 4.2 program [35–37] using our default settings of 25°C, 3 mM MgCl_2_ and 50mM NaCl. The stability of palindrome structures was routinely quantified in the presence of short flanking single strand regions for uniform alignment, although the listed coordinates exclude these tails. As a control, we characterized the palindromes at the junctions of inverted repeats formed in a yeast model system [7]. The UNAFold® software (RPI 2017-042) is under license 80-00840 from Rensselaer Polytechnic Institute. We also analyzed the potential palindromes in the 200 bp of genomic sequence surrounding the incipient site of recombination in TRP/INVs exhibiting deletions of ≤ 40 bp.

The UNAfold algorithm most often generated simple hairpin predictions. However, three types of other pairing patterns were also observed. Structures in which the hairpin was closely adjacent to more extensive base pairing were termed extended forms, while predicted patterns containing two or more adjacent hairpins were termed multi-palindromic. Structures that were paired but appeared to contain some form of distortion were termed complex forms. Examples of these forms are shown in SI Fig. 4.

Overall palindrome predictions for centromeric and non-centromeric were determined by Palindrome Analyzer [39], using parameters exhibited by the majority of palindromes associated with the TRP/INVs from centromeric or dispersed domains of chromosomes. The same parameters were then used for the entire genome. We first determined that the most frequent classes of the terminal portions of centromere 3 TRP/INV-associated palindromes. This class had stem sizes of 12-100 bp, loop sizes of 4-8 bp, with 0-2 stem mismatches. We then identified these partial structures in the output of the Palindrome Analyzer [39] and analyzed them for the presence of part or all of the input palindrome structure. These ΔG_cf_-ΔG_lin_ values generally ranged from 10-16 kcal/mol. Since the distribution was similar for the observed and predicted palindromes, we determined the number of predicted palindromes as the sum of all candidates within this range. However, this number was corrected for that structures that had stabilities lower than the study threshold. For each centromere, we added candidates in the 10-16 kcal/mole range and adjusted these for the expected number of false positives.

We could not apply to non-centromeric sequence due to the differing conditions identified in TRP/INV associated in palindromes in AT-rich and dispersed regions. Examination non-centromeric palindromes from chromosomes 1, 3 and 4 revealed stem lengths ranging from 5-12 bp, loops ranging from 4-19 bp with stem mismatches from 0-1. We empirically converted ΔG into ΔG_cf_-ΔG_lin_ values by first remeasuring ΔG at 37°C, 50mM NaCl, and 1mM MgCl_2.,_ The values, representing the stability of a hairpin, were then doubled to generate ΔG_cf_-ΔG_lin_, a measure of cruciform stability. ΔG_cf_-ΔG_lin_ values ranging from 3-12 kcal/mol distributed as follows: 15 at 3-6 kcal/mol (Class 1), 10 at 6.1-9 kcal/mol (Class 2) and 14 from at kcal/mol (Class 3). Since the predicted output from Palindrome Analyzer for a 60 kb region of centromere 3 had a different distribution (24 in class 1, 105 in class 2, and 507 in class 3) than the TRP/INV-associated palindromes, we normalized the output values using a weighted eligibility model [40, 41]. Specifically, to estimate how many palindrome sites in a genomic interval are actually competent to initiate recombination, we compared the observed distribution across palindrome classes with the relative abundance of candidates. For each class 𝑖, we calculated a per-candidate usage rate 𝑟_𝑖_ = 𝑈_𝑖_/𝑛_𝑖_, where 𝑈_𝑖_ is the number of observed TRP/INV-associated palindromes and 𝑛_𝑖_ is the number of candidate palindromes in the interval. To account for the unequal usage across classes, we normalized by the maximum observed rate, defining a relative eligibility fraction ℎ_𝑖_ = 𝑟_𝑖_/max _𝑗_ 𝑟_𝑗_. The estimated pool of competent candidates in the interval is described by

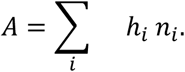

### Sequence Structure in Centromeric and Subtelomeric ATEs

To ascertain the frequency of short repeats within ATEs, we identified 250 bp DNAs from centromeric and subtelomeric ATEs having a 90% A+T content and assayed for self-homology by BLAST analysis to identify direct, inverted, and palindromic repeats. We rated ungapped regions of homology as a function of the BLAST e-value. We compared this distribution with 250 bp sequences derived from an Excel random sequence generator (Microsoft) set at 90%A+T DNA. Both perfect and mismatched repeats at each value were combined for the analysis of direct and inverted repeats. To quantify the statistical significance of the deviation from nonrandom behavior, we evaluated the number of sequences containing at least one event present at e-values of <10^-4^ for genomic and random datasets by the Fisher Exact Test. Perfectly matched palindromes were similarly analyzed at BLAST e values of <10^-8^ and evaluated by the Fisher Exact Test.

## DATA ACCESS

The raw data from nanopore and PacBio sequencing are publicly available at NCBI SRA listed as BioProject PRJNA1193636 with the following BioSample accession numbers: SAMN45150363 (G0 Nanopore), SAMN45150364 (G5 Nanopore), SAMN45150365 (G10 Nanopore), SAMN45150366 (G15 Nanopore), SAMN45150367 (G15 PacBio-1), SAMN45150368 (G15 PacBio-2). The G0 and G15 assemblies are present in the Sequence Cloud Server and are available on request.

## ACKNOWLEDGEMENTS

We would like to thank and Bonnie Brewer (U of Washington) Drs. Prescott Deininger (Tulane), Matthew Lebar (USDA), Brian Mack (USDA), and Bill Wimley (Tulane), for stimulating discussions during these studies. We would also like to give our appreciation to Melody Baddoo and for bioinformatic assistance. Drs. Mary J Clancy (UNO) and Bibo Li (Cleveland State University), and Bonnie Hoffman are thanked for critical reading of the manuscript.

## Author contributions

Conceptualization: AJL

Methodology: AJL, JWC

Investigation: JWC, AM, AS, RL, AJL

Visualization: AJL, AS

Supervision: AJL, EF

Writing—original draft: AJL

Writing—review & editing: AJL, JWC, EF, AM, AS

## Data and materials availability

All data presented herein is available to the research community.

