## Supplemental Information for "Clustering of Inverted Triplications in Centromeric and Subtelomeric Chromosomal Regions of *Aspergillus flavus*"

Jeffrey W. Cary *et al*.

**Table of Contents:**

| | | |
|---|---|---|
| SI _Fig_S1 | Fine Structure Analysis of Suspected Heritable TRP/INVs | **2** |
| SI_Fig_S2 A, B | BLAST Assay for Non-ATE TRP/INVs. | **3-4** |
| SI _Fig_S2: C, D | BLAST Assay for Subtelomeric and Centromeric TRP/INVs. | **5-6** |
| SI_Fig_S3 A, B | BLAST Assay for Palindromic Sequence from Non-Centromeric and Subtelomeric TRP/INVs. | **7-8** |
| _Fig_S3 C, D | BLAST Assay for Palindromic Sequence from Centromeric and Subtelomeric TRP/INVs. | **9-10** |
| SI_Fig_S4 | Examples of Extended and Complex Structures | **11** |
| SI_Table_S1 | | **11-18** |
| SI_Table_S2 | | **19** |
| SI_Table_S3A | | **20-21** |
| SI_Table_S3B | | **22-26-** |
| SI_Table_S3C | | **27** |

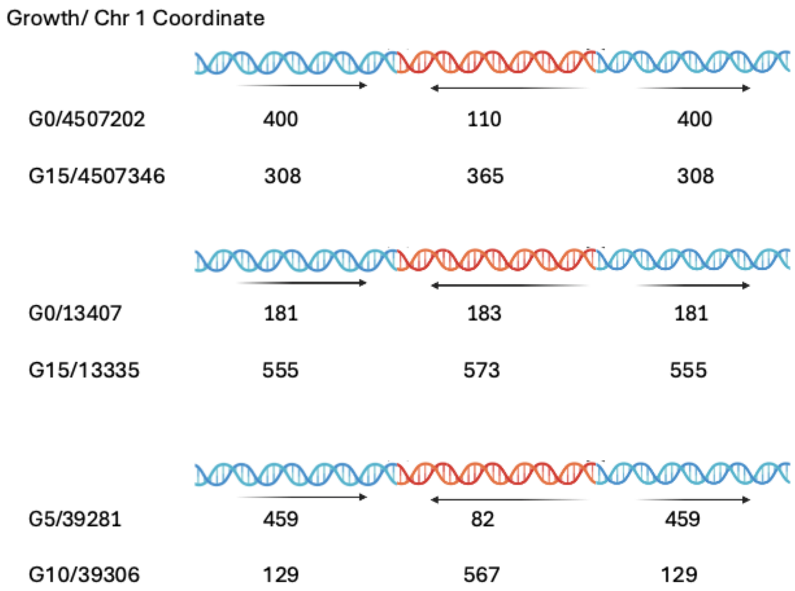

SI_Fig_S1 **Fine Structure Analysis of Suspected Heritable TRP/INVs.**

The TRP/INVs present in Chromosome 1 (Chr1)from G5, G10 and G15 cultures were compared to one another. Three candidates suspected to be at the same chromosome coordinates based on the Sniffles 2 were analyzed by BLAST to the genomic assembly. The structure of each TRP/INV is shown together with the culture stage (G5-G15) and Chr1 coordinate. The general structure of the TRP/INV is listed above each cluster as described in Figure 1.

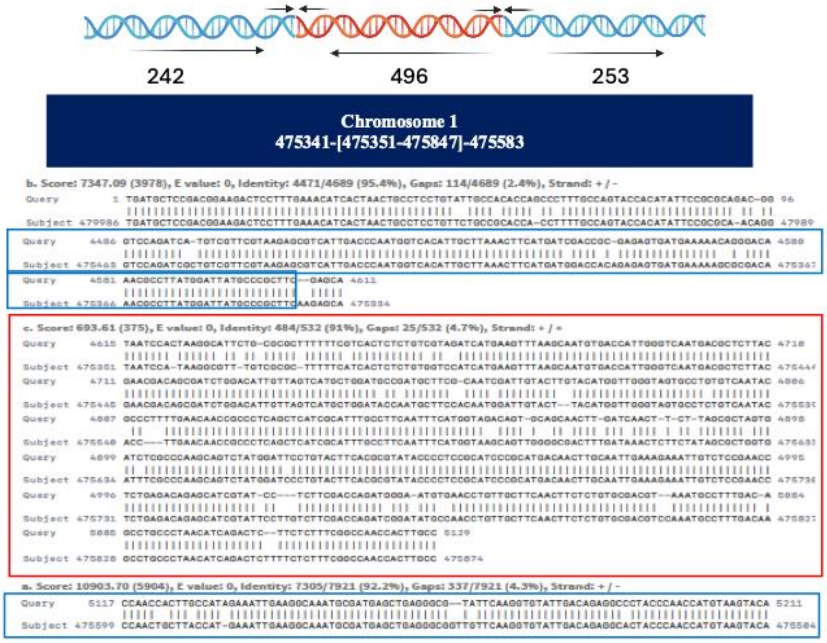

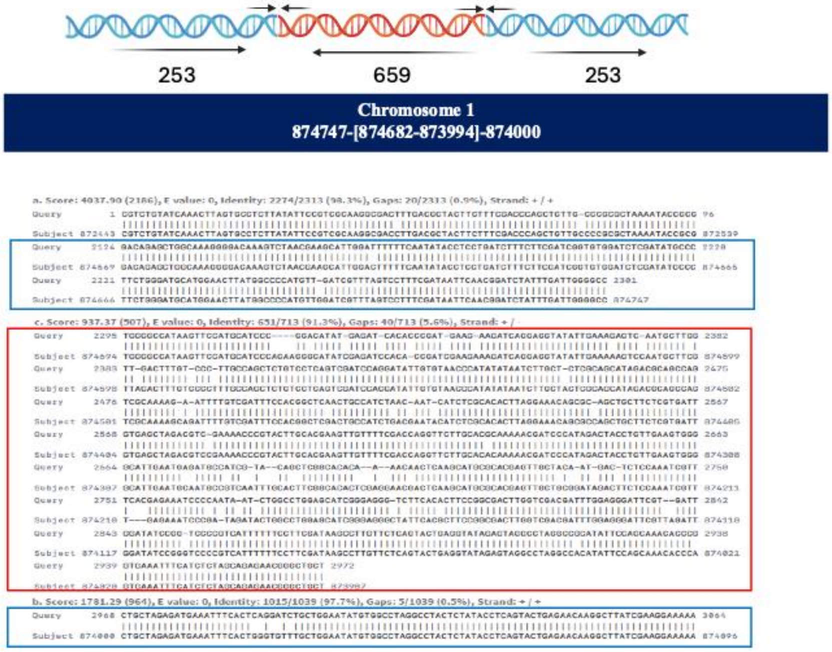

SI_Fig_S2A, B. **BLAST Assay for Non-ATE TRP/INVs.**

The junctional structures of the two independents TRP/INVs from dispersed non-ATE regions on chromosome were determined by BLAST analysis and the length of direct (blue) and inverted (red) repeats were calculated. The detailed coordinates of direct and inverted (bracketed) sequences are shown. The BLAST reads of the left junction and right junctions are shown together with the statistical output of the BLAST results from SequenceServer. +/- refers to a sequence complementary to the query sequence, while +/+ refers to a sequence in same orientation as the query sequence. The query is the read sequence, and the subject is the G15 assembly. Blue and red boxes refer to the duplicated and inverted sequences, respectively.

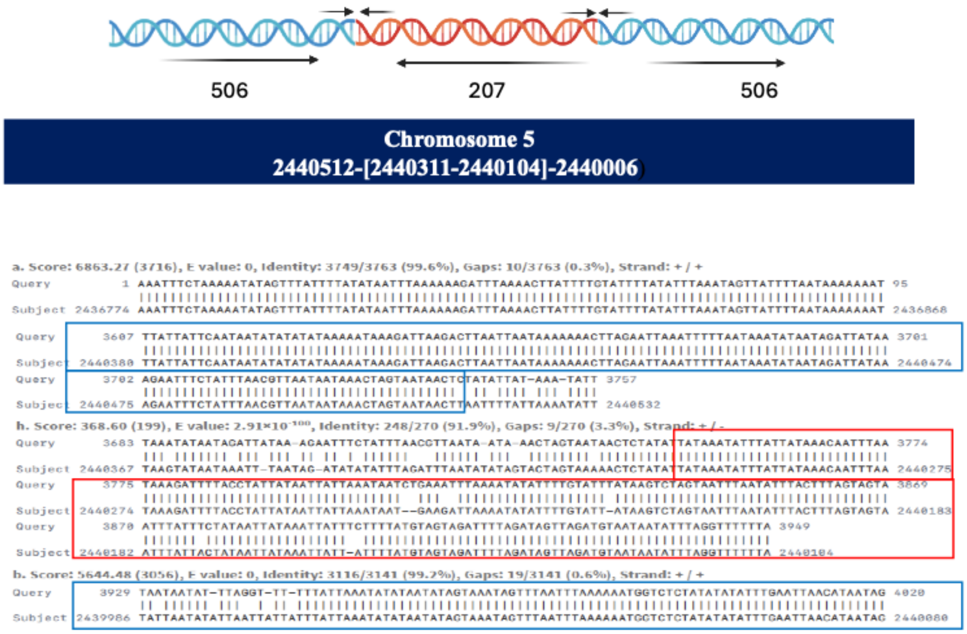

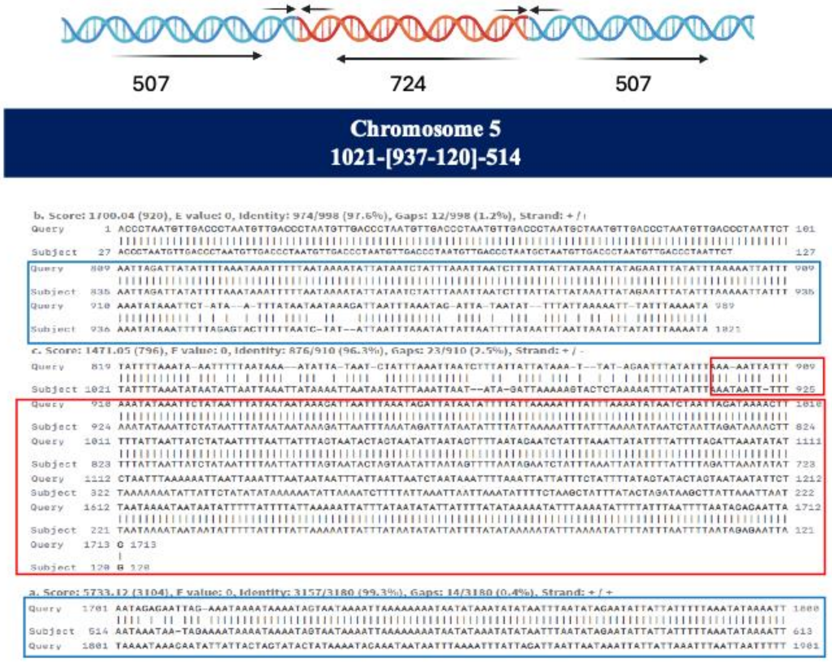

SI_Fig_S2C,D. **BLAST Assay for Subtelomeric and Centromeric TRP/INVs**

BLAST analysis was used to determine both the junctional structures of the two independent

TRP/INVs from clustered centromeric (C) and subtelomeric (D) regions on chromosome 5 and the length of direct (blue) and inverted (red) repeat. The detailed coordinates of direct and inverted (bracketed) are shown. The BLAST reads were characterized as described in SI_Fig_S2A.

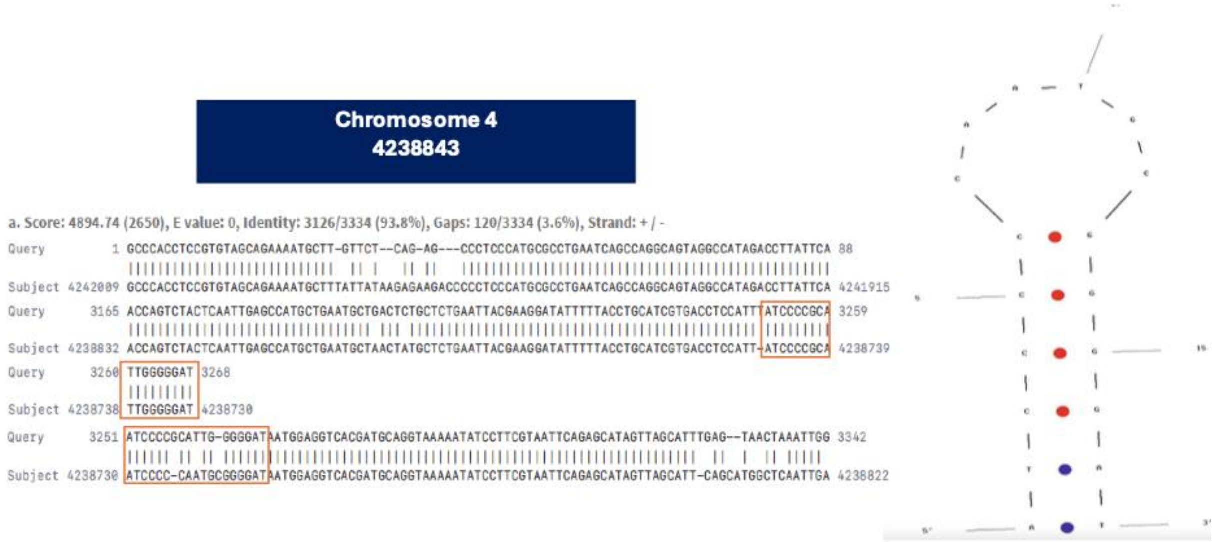

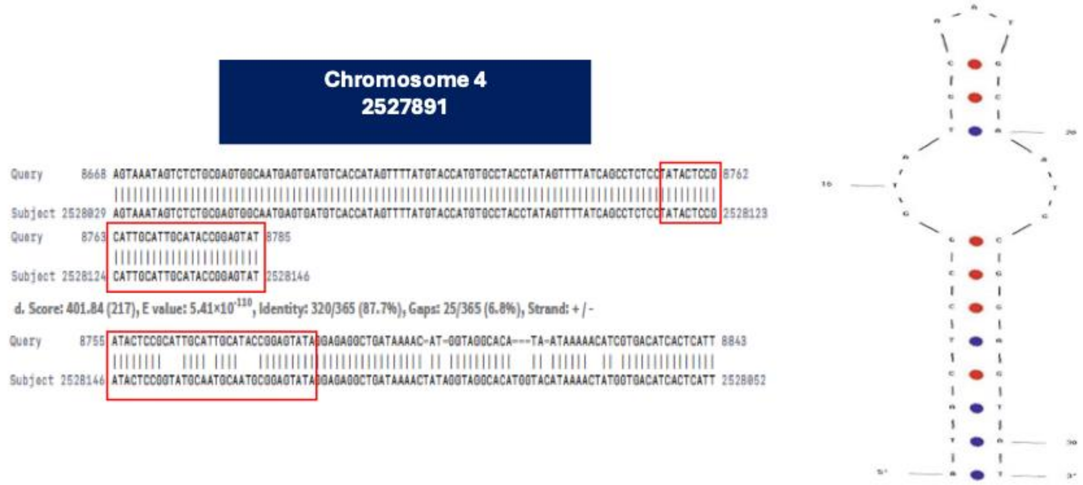

SI_Fig_S3A, B**. BLAST Assay for Palindromic Sequence from Non-Centromeric and Subtelomeric TRP/INVs.**

Palindromes determined by BLAST analysis showing the junctional structures of two independent TRP/INVs from dispersed non-ATE regions on chromosomes 4 at the designated coordinates. Red boxes shown the palindromic sequences. The putative structure determined by UNAfold is shown on the right with red dots and blue dots referring to G-C and A-T base pairs, respectively.

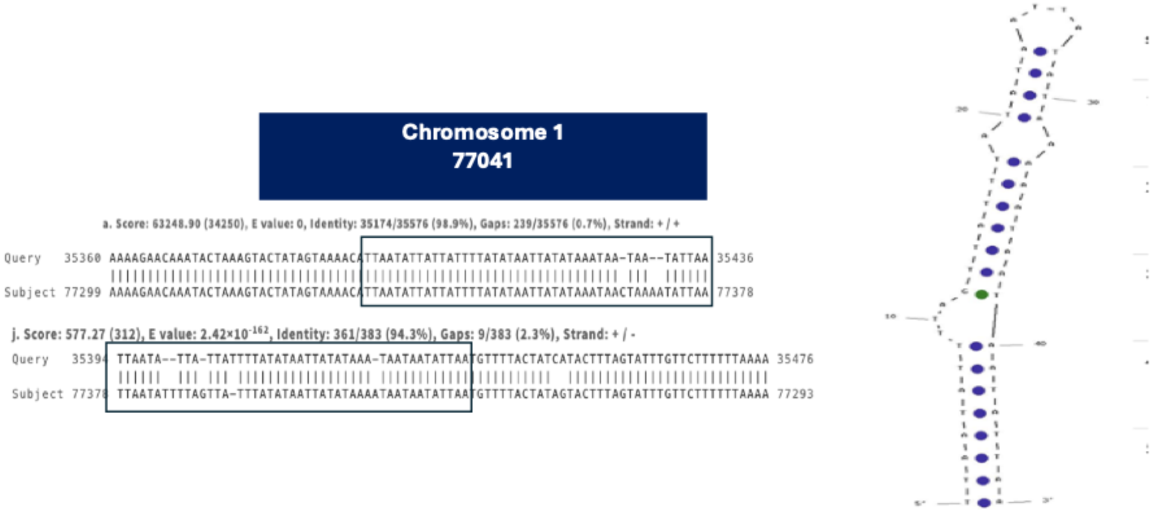

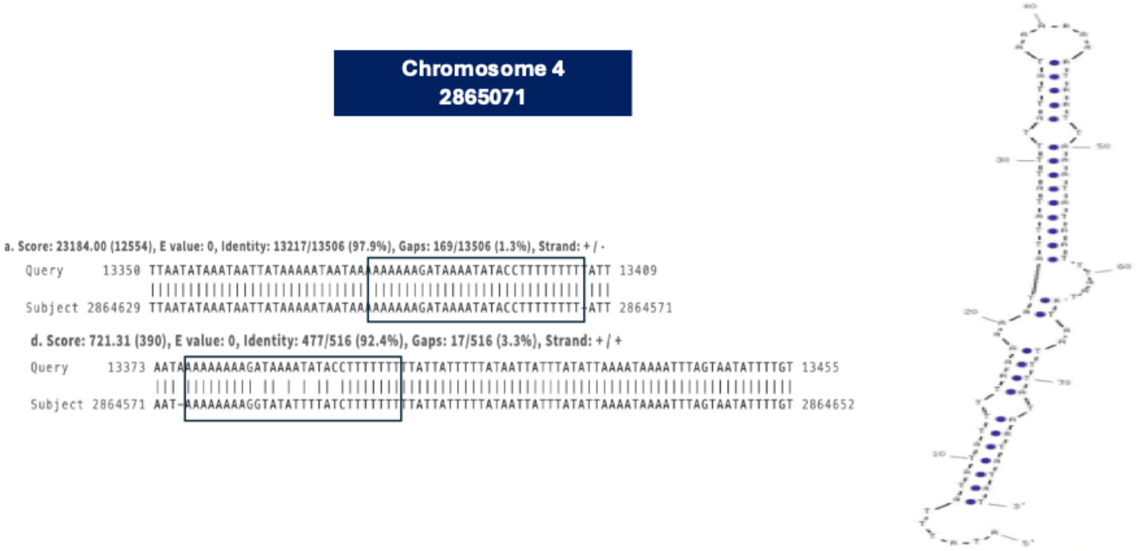

SI_Fig_S3 C,D. **BLAST Assay for Palindromic Sequence from Centromeric and Subtelomeric TRP/INVs.**

Palindromes determined by BLAST analysis showing the junctional structure of the two independent TRP/INVs from ATE regions on chromosomes 1 and 4. Red boxes show the palindromic sequences. The putative structure determined by UNAfold is shown on the right with blue dots referring to A-T base pairs.

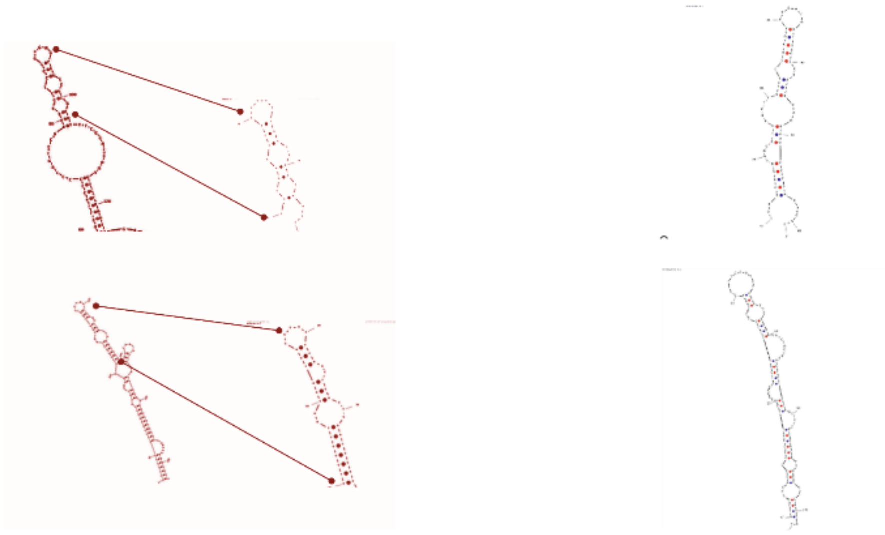

A. Extended Forms B. Complex Forms

SI Figure_4: **Examples of Extended and Complex Pairing Patterns Associated with TRP/INVs Predicted from UNAfold.**

(A, Extended Forms ) Two examples of the extended pairings (left) adjacent to the structure used for the thermodynamic stability calculations (right). (top) The structure formed on chromosome 2, 4171485-4171510 and (bottom) the structure formed at chromosome 3, 1230017-1230054 as described in Supplementary Table 3B. (B, Complex Forms) Two examples of unusual pairing patterns identified in a subset of regions. (top) The structure predicted from chromosome 5, 1759115-1759167 and (bottom) the structure predicted from chromosome 6, 561314-561352 as described in SI Table 3B.

| Chr | coordinate | direct repeat (bp) | inverted repeat (bp) | Deletions Left (bp) | Deletions Right (bp) | TID Type | Gene (F9C07#) |
|---|---|---|---|---|---|---|---|
| 1 | 13335 | 555 | 573 | 37 | 43 | 1 | NC |
| 1 | 77041 | 329 | 329 | 0 | 20 | 1 | NC |
| 1 | 475416 | 242 | 496 | 1 | 264 | 2 | 2134323 |
| 1 | 874229 | 253 | 312 | 65 | 16 | 2 | 2278780 |
| 1 | 937833 | 249 | 312 | 19 | 26 | 1 | 1015060 |
| 1 | 1270243 | 131 | 270 | 116 | 83 | 3 | 2135501 |
| 1 | 1565531 | 208 | 102 | 134 | 42 | 2 | 2278999 |
| 1 | 2265912 | 192 | 143 | 81 | 0 | 2 | 834 |
| 1 | 2286434 | 400 | 462 | 0 | 58 | 2 | 1055498 |
| 1 | 2300803 | 484 | 306 | 169 | 9 | 2 | 848 |
| 1 | 3240299 | 172 | 464 | 16 | 298 | 1 | 2228016 |
| 1 | 3436109 | 307 | 369 | 26 | 7 | 1 | NC |
| 1 | 3485461 | 140 | 249 | 76 | 11 | 2 | 2228016 |
| 1 | 3562082 | 1237 | 1211 | 23 | 3 | 1 | 1310 |
| 1 | 3750785 | 572 | 500 | 50 | 18 | 1 | NC |
| 1 | 3893569 | 531 | 461 | 90 | 20 | 2 | 1448 |
| 1 | 4140799 | 419 | 415 | 16 | 17 | 1 | NC |
| 1 | 4449595 | 187 | 193 | 86 | 76 | 3 | 1647 |
| 1 | 4459765 | 276 | 252 | 41 | 22 | 1 | NC |
| 1 | 4507346 | 308 | 365 | 10 | 84 | 2 | NC |
| 1 | 4510493 | 254 | 233 | 39 | 0 | 1 | NC |
| 1 | 4514526 | 158 | 174 | 36 | 19 | 1 | NC |
| 1 | 4519524 | 642 | 603 | 56 | 16 | 2 | NC |
| 1 | 4528244 | 743 | 149 | 518 | 76 | 3 | NC |
| 1 | 4529167 | 245 | 335 | 35 | 97 | 2 | NC |
| 1 | 4538202 | 190 | 598 | 19 | 321 | 2 | NC |
| 1 | 4539603 | 141 | 345 | 26 | 230 | 2 | NC |
| 1 | 4603128 | 590 | 271 | 4 | 325 | 2 | NC |
| 1 | 4604416 | 526 | 97 | 51 | 820 | 3 | NC |
| 1 | 4615935 | 138 | 333 | 0 | 253 | 2 | NC |
| 1 | 4648368 | 185 | 223 | 20 | 27 | 1 | 2228454 |
| 1 | 4868805 | 296 | 316 | 32 | 42 | 1 | ND |
| 1 | 4988696 | 203 | 254 | 50 | 19 | 1 | 2279968 |
| 1 | 5016740 | 361 | 373 | 0 | 16 | 1 | 2279977 |

| | | | | | | | |
|---|---|---|---|---|---|---|---|
| 2 | 409525 | 165 | 375 | 26 | 184 | 2 | NC |
| 2 | 454494 | 160 | 182 | 0 | 0 | 1 | NC |
| 2 | 493401 | 238 | 348 | 0 | 133 | 2 | 2197617 |
| 2 | 1804843 | 313 | 360 | 40 | 87 | 2 | 2277186 |
| 2 | 2064565 | 246 | 265 | 0 | 22 | 1 | NC |
| 2 | 2106651 | 863 | 163 | 646 | 54 | 3 | NC |
| 2 | 2106651.1 | 78 | 261 | 64 | 125 | 3 | NC |
| 2 | *2121580* | 787 | 800 | 10 | 27 | 1 | NC |
| 2 | 2303657 | 2125 | 2011 | 58 | 11 | 2 | 1216492 |
| 2 | 2600903 | 554 | 460 | 118 | 20 | 2 | 2128422 |
| 2 | 2674476 | 321 | 313 | 30 | 86 | 2 | NC |
| 2 | 3246057 | 197 | 276 | 33 | 101 | 2 | 2277572 |
| 2 | 3348375 | 787 | 723 | 50 | 4 | 2 | 2225896 |
| 2 | 3365195 | 382 | 408 | 0 | 12 | 1 | 2217776 |
| 2 | 3673467 | 373 | 378 | 54 | 23 | 2 | NC |
| 2 | 3840594 | 542 | 555 | 0 | 35 | 1 | 2217863 |
| 2 | 3895441 | 1429 | 1199 | 231 | 15 | 2 | 2277774 |
| 2 | 4115827 | 502 | 550 | 131 | 0 | 2 | NC |
| 2 | 4172335 | 1042 | 859 | 190 | 4 | 2 | NC |
| 2 | 4274226 | 321 | 351 | 0 | 57 | 2 | 2402 |
| 2 | 5407930 | 240 | 240 | 0 | 65 | 2 | 2278219 |
| 2 | 5551182 | 308 | 301 | 40 | 17 | 1 | 2056731 |
| 2 | 6204471 | 2365 | 1340 | 50 | 58 | 2 | 1356924 |
| 3 | 268975 | 316 | 238 | 13 | 8 | 1 | 2230157 |
| 3 | 567261 | 282 | 83 | 0 | 186 | 2 | 4362 |
| 3 | 572934 | 1350 | 1266 | 10 | 102 | 2 | 4364 |
| 3 | 615936 | 303 | 382 | 0 | 79 | 2 | NC |
| 3 | 952213 | 139 | 311 | 48 | 237 | 3 | 2230400 |
| 3 | 962141 | 531 | 342 | 188 | 27 | 3 | NC |
| 3 | 1064285 | 310 | 373 | 7 | 77 | 2 | 2104432 |
| 3 | 1230011 | 188 | 187 | 21 | 37 | 1 | NC |
| 3 | 1526908 | 218 | 450 | 0 | 232 | 2 | 2281754 |
| 3 | 1572690 | 700 | 707 | 27 | 0 | 1 | NC |
| 3 | 1734385 | 653 | 241 | 645 | 0 | 2 | NC |
| 3 | 2424515 | 181 | 388 | 10 | 215 | 2 | 4790 |
| 3 | 2455821 | 181 | 189 | 12 | 17 | 1 | NC |

| | | | | | | | |
|---|---|---|---|---|---|---|---|
| 3 | 2531871 | 591 | 909 | 21 | 646 | 2 | NC |
| 3 | 2693002 | 216 | 419 | 0 | 203 | 2 | NC |
| 3 | 2705388 | 247 | 277 | 0 | 42 | 1 | NC |
| 3 | 2730296 | 545 | 154 | 367 | 24 | 2 | NC |
| 3 | 2732735 | 530 | 850 | 0 | 20 | 1 | NC |
| 3 | 2737298 | 234 | 126 | 111 | 0 | 2 | NC |
| 3 | 2739469 | 191 | 217 | 0 | 16 | 1 | NC |
| 3 | 2739747 | 589 | 90 | 60 | 435 | 3 | NC |
| 3 | 2741124 | 686 | 693 | 24 | 24 | 1 | NC |
| 3 | 2750522 | 690 | 171 | 66 | 454 | 3 | NC |
| 3 | 2762452 | 154 | 190 | 0 | 0 | 1 | NC |
| 3 | 2787461 | 209 | 66 | 163 | 19 | 2 | NC |
| 3 | 2789492 | 536 | 814 | 33 | 305 | 2 | NC |
| 3 | 2790068 | 574 | 675 | 4 | 62 | 1 | NC |
| 3 | 2812701 | 537 | 432 | 48 | 72 | 2 | NC |
| 3 | *2844927* | 357 | 342 | 15 | 0 | 1 | NC |
| 3 | 2868799 | 381 | 440 | 63 | 105 | 3 | 1428495 |
| 3 | 2922743 | 303 | 278 | 10 | 23 | 1 | 2066186 |
| 3 | 3287705 | 379 | 421 | 0 | 20 | 1 | 1439314 |
| 3 | 3293049 | 89 | 327 | 27 | 257 | 2 | 7152 |
| 3 | 3792774 | 497 | 497 | 34 | 275 | 1 | 7336 |
| 3 | 3837633 | 203 | 26 | 24 | 0 | 1 | 1456142 |
| 3 | 3837633.2 | 194 | 201 | 31 | 25 | 1 | 1456142 |
| 3 | 3935716 | 7269 | 7459 | 102 | 598 | 3 | 2282455 |
| 3 | 4213435 | 195 | 581 | 16 | 407 | 2 | 2282540 |
| 3 | 4213435.2 | 106 | 346 | 34 | 275 | 1 | 2282540 |
| 3 | 4712346 | 400 | 412 | 25 | 25 | 1 | 2282685 |
| 3 | 4939779 | 387 | 406 | 0 | 27 | 1 | 2282762 |
| 3 | 4990724 | 239 | 195 | 37 | 2 | 1 | 2255881 |
| 3 | 5171690 | 292 | 238 | 116 | 32 | 2 | NC |
| 3 | 5192156 | 957 | 74 | 851 | 6 | 2 | NC |
| 4 | 2545 | 1507 | 1465 | 40 | 0 | 1 | NC |
| 4 | 5473 | 214 | 98 | 36 | 81 | 2 | NC |
| 4 | 43041 | 130 | 263 | 19 | 145 | 2 | 2282857 |
| 4 | 568528 | 153 | 155 | 0 | 7 | 1 | 3373;<br>2283015 |
| 4 | 694069 | 408 | 490 | 8 | 95 | 2 | t173 |

| | | | | | | | |
|---|---|---|---|---|---|---|---|
| 4 | 794961 | 440 | 496 | 0 | 66 | 1 | 2283081 |
| 4 | 870310 | 868 | 958 | 30 | 117 | 2 | 3478 |
| 4 | 1063346 | 277 | 274 | 10 | 2 | 1 | NC |
| 4 | 1194513 | 293 | 492 | 0 | 10 | 1 | 3605 |
| 4 | 1273134 | 123 | 299 | 0 | 184 | 2 | 2283226 |
| 4 | 1275195 | 182 | 189 | 15 | 0 | 1 | 2283226 |
| 4 | 1607156 | 361 | 326 | 8 | 15 | 1 | 2283316 |
| 4 | 2308858 | 335 | 174 | 104 | 71 | 3 | NC |
| 4 | 2527891 | 317 | 362 | 0 | 52 | 2 | NC |
| 4 | 2542570 | 198 | 276 | 74 | 148 | 3 | NC |
| 4 | 2568055 | 290 | 540 | 37 | 288 | 2 | NC |
| 4 | 2818293 | 321 | 245 | 0 | 34 | 1 | NC |
| 4 | 2821444 | 923 | 905 | 125 | 116 | 3 | NC |
| 4 | 2861376 | 555 | 261 | 43 | 251 | 2 | NC |
| 4 | 2865070 | 447 | 505 | 42 | 0 | 1 | NC |
| 4 | 3150384 | 219 | 241 | 12 | 40 | 1 | NC |
| 4 | 3431318 | 244 | 290 | 8 | 64 | 2 | 2159007 |
| 4 | 3869842 | 285 | 368 | 29 | 121 | 2 | 2283931 |
| 4 | 4238843 | 366 | 353 | 0 | 0 | 1 | 2284036 |
| 4 | 4239136 | 1156 | 1065 | 124 | 5 | 2 | 2284036 |
| 4 | 4414808 | 321 | 269 | 16 | 0 | 1 | 2284092 |
| 4 | 4623095 | 420 | 502 | 31 | 110 | 2 | 1603332 |
| 5 | 527 | 507 | 745 | 0 | 394 | 2 | NC |
| 5 | 1790 | 525 | 497 | 0 | 0 | 1 | NC |
| 5 | 4047 | 200 | 142 | 60 | 27 | 2 | NC |
| 5 | 346121 | 319 | 329 | 24 | 33 | 1 | 2284336 |
| 5 | 376113 | 835 | 797 | 57 | 5 | 2 | 6834 |
| 5 | 495710 | 198 | 414 | 26 | 257 | 2 | NC |
| 5 | 618809 | 296 | 377 | 0 | 80 | 2 | NC |
| 5 | 636592 | 217 | 136 | 54 | 17 | 2 | NC |
| 5 | 1566481 | 276 | 318 | 8 | 46 | 1 | 2284739 |
| 5 | 1706097 | 58 | 374 | 14 | 336 | 2 | 2284739 |
| 5 | 1759989 | 1154 | 1139 | 39 | 7 | 1 | 6308 |
| 5 | 1847949 | 278 | 149 | 101 | 16 | 2 | NC |
| 5 | 2017485 | 518 | 519 | 25 | 25 | 1 | 6206 |
| 5 | 2219537 | 254 | 208 | 196 | 150 | 3 | 6134 |
| 5 | *2313831* | 266 | 231 | 39 | 31 | 1 | 6102 |

| | | | | | | | |
|---|---|---|---|---|---|---|---|
| 5 | 2332008 | 203 | 328 | 0 | 88 | 2 | NC |
| 5 | 2406521 | 356 | 544 | 13 | 201 | 2 | NC |
| 5 | 2431186 | 735 | 59 | 45 | 269 | 2 | NC |
| 5 | 2432481 | 180 | 210 | 12 | 42 | 1 | NC |
| 5 | 2440449 | 506 | 207 | 201 | 98 | 3 | NC |
| 5 | 2447902 | 492 | 848 | 0 | 356 | 2 | NC |
| 5 | 2457207 | 257 | 653 | 19 | 435 | 2 | NC |
| 5 | 2472764 | 235 | 586 | 0 | 51 | 2 | NC |
| 5 | 2482236 | 446 | 717 | 0 | 271 | 2 | NC |
| 5 | 2485809 | 206 | 339 | 4 | 132 | 2 | NC |
| 5 | *2813438* | 197 | 319 | 11 | 133 | 2 | 2093803 |
| 5 | 3388929 | 437 | 413 | 69 | 54 | 3 | 2234619 |
| 5 | 3446394 | 1529 | 1583 | 0 | 47 | 1 | 2245974 |
| 5 | 3772497 | 353 | 314 | 28 | 0 | 1 | 9062 |
| 5 | 4111924 | 161 | 235 | 0 | 74 | 2 | *ustD* |
| 5 | 4258504 | 637 | 766 | 21 | 149 | 2 | 2071928 |
| 5 | 4510858 | 1067 | 1032 | 12 | 0 | 1 | NC |
| 5 | 4543604 | 674 | 50 | 500 | 14 | 2 | NC |
| 5 | 4547799 | 643 | 643 | 47 | 43 | 1 | NC |
| 6 | 561512 | 304 | 298 | 39 | 11 | 1 | 2285823 |
| 6 | 672926 | 197 | 485 | 19 | 330 | 2 | *dit2* |
| 6 | 1167658 | 291 | 308 | 0 | 20 | 1 | 2286014 |
| 6 | 1257940 | 188 | 187 | 41 | 43 | 1 | 8308 |
| 6 | 1298630 | 536 | 538 | 48 | 48 | 1 | NC |
| 6 | 1661374 | 164 | 189 | 0 | 38 | 1 | 2246838 |
| 6 | 2149517 | 312 | 264 | 22 | 0 | 1 | NC |
| 6 | 2204802 | 372 | 98 | 27 | 250 | 2 | NC |
| 6 | 2486587 | 248 | 248 | 0 | 0 | 1 | 12042 |
| 6 | 2523037 | 410 | 454 | 0 | 94 | 1 | NC |
| 6 | 2702072 | 165 | 304 | 21 | 157 | 2 | 2286450 |
| 6 | 2745519 | 230 | 273 | 24 | 57 | 2 | 1799237 |
| 6 | 2824544 | 175 | 151 | 67 | 45 | 2 | NC |
| 6 | 3622215 | 123 | 225 | 51 | 145 | 2 | NC |
| 7 | 89355 | 144 | 223 | 35 | 104 | 2 | 5171 |
| 7 | 109150 | 593 | 575 | 35 | 7 | 1 | NC |

| | | | | | | | |
|---|---|---|---|---|---|---|---|
| 7 | 191198 | 306 | 239 | 105 | 19 | 2 | NC |
| 7 | 289804 | 414 | 535 | 14 | 132 | 2 | 5204 |
| 7 | 416095 | 223 | 194 | 80 | 38 | 2 | NC |
| 7 | 448368 | 338 | 76 | 41 | 218 | 2 | 5306 |
| 7 | 690328 | 170 | 157 | 0 | 17 | 1 | NC |
| 7 | 1169343 | 873 | 900 | 29 | 57 | 2 | 2280776 |
| 7 | 1207830 | 579 | 677 | 47 | 136 | 2 | 1849043 |
| 7 | 1656240 | 116 | 370 | 37 | 291 | 2 | 2145462 |
| 7 | 1712436 | 263 | 246 | 154 | 137 | 3 | 5787 |
| 7 | 1764947 | 375 | 379 | 21 | 67 | 2 | 1866164 |
| 7 | 1815039 | 467 | 231 | 236 | 0 | 2 | 2200969 |
| 7 | 1865225 | 524 | 549 | 18 | 22 | 1 | NC |
| 7 | 2302557 | 4066 | 5244 | 19 | 1219 | 2 | 1883257 |
| 7 | 2575541 | 381 | 415 | 15 | 49 | 1 | NC |
| 7 | 2617758 | 196 | 184 | 18 | 6 | 1 | NC |
| 7 | 2618846 | 586 | 712 | 0 | 20 | 1 | NC |
| 7 | 2630527 | 238 | 208 | 0 | 24 | 1 | NC |
| 7 | 2646144 | 526 | 611 | 0 | 0 | 1 | NC |
| 8 | 256862 | 465 | 253 | 0 | 212 | 2 | NC |
| 8 | 731953 | 1310 | 1543 | 28 | 257 | 2 | 2278731 |
| 8 | 797525 | 341 | 36 | 287 | 17 | 2 | 2278750 |
| 8 | 809155 | 178 | 189 | 6 | 27 | 1 | NC |
| 8 | 841463 | 202 | 155 | 60 | 10 | 2 | 2134869 |
| 8 | 919960 | 449 | 498 | 20 | 61 | 2 | 2134952 |
| 8 | 1428248 | 360 | 344 | 31 | 0 | 1 | NC |
| 8 | 1769900 | 291 | 284 | 11 | 7 | 1 | 665 |
| 8 | 1814495 | 384 | 490 | 0 | 136 | 2 | NC |
| 8 | 1825065 | 310 | 332 | 19 | 41 | 1 | NC |
| 8 | 1852346 | 639 | 683 | 0 | 37 | 1 | NC |
| 8 | 1869428 | 621 | 790 | 21 | 197 | 2 | NC |
| 8 | 1869802 | 355 | 548 | 0 | 229 | 1 | NC |
| 8 | 1896688 | 342 | 657 | 0 | 313 | 2 | NC |
| 8 | 1898604 | 343 | 340 | 25 | 22 | 1 | NC |
| 8 | 1905303 | 631 | 194 | 460 | 24 | 2 | NC |
| 8 | 2160564 | 473 | 794 | 37 | 639 | 2 | NC |
| 8 | 2314298 | 107 | 233 | 10 | 136 | 2 | 853 |
| 8 | 2319292 | 112 | 207 | 5 | 104 | 2 | 2279203 |

| | | | | | | | |
|---|---|---|---|---|---|---|---|
| 8 | *2580536* | 201 | 203 | 29 | 27 | 1 | NC |
| 8 | 2581800 | 63 | 308 | 20 | 280 | 2 | NC |
| 8 | 2738339 | 219 | 293 | 22 | 3 | 1 | 2279334 |
| 8 | 2804414 | 62 | 242 | 21 | 257 | 2 | NC |
| 8 | 3239040 | 253 | 292 | 0 | 57 | 2 | NC |
| 8 | 3241336 | 137 | 247 | 36 | 146 | 2 | NC |

SI_**Table**_1. **Characteristics of G15 TRP/INVs Identified by ONT Sequencing.** Quantitative analysis was performed with variant read values (DV) of 1 as described in Materials and Methods. The chromosomal sites (Chr) are shown listed together with the size of the TRP/INV direct and inverted repeats and the approximate amount of approximate junctional sequence lost (black) or present in a palindrome (red) in the chromosomal DNA. Based on this analysis the TRP/INV is then placed into Type 1 (symmetric, both junctions having a gap of ≤ 50 bp), Type 2 (one junction having a gap of >50 bp) or Type 3 (two junctions having a gap of >50 bp). The right most column designates the coding [R9C07# (*1*) or gene designation (in italics)] or non-coding (NC) location of the TRP/INV. The yellow color refers to a centromeric TRP/INV, while the orange color denotes a TRP/INV within subtelomeric or other ATE sequences. Uncolored boxes refer to TRP/INVs located at other genomic sites. The .1 designation indicates two independent TRP/INVs at the coordinate. Italics indicate samples were not designated in a domain due to their ambiguous position.

| Chr | Coordinate | Direct Repeat (bp) | Inverted Repeat (bp) | pTID Type |
|---|---|---|---|---|
| 2 | 2063917 | 158 | 300 | 2 |
| 3 | 2749349 | 180 | 101 | 2 |
| 3 | 2824589 | 180 | 53 | 2 |
| 5 | 2492048 | 200 | 424 | 2 |
| 5 | 4544691 | 206 | 309 | 2 |
| 6 | 2132490 | 177 | 46 | 2 |
| 8 | 1874224 | 420 | 289 | 2 |

SI_Table_S2. **Characteristics of TRP/INVs Identified by PacBio Sequencing**. Quantitative identification of TRP/INVs from PacBio sequenced DNA was performed **as** as described in Table S1.

| Chr | Deletion | Palindrome (start) | Location (stop) | Length (bp) | ΔG ( kcal/mol) | Tm (°C) | Paired Extensions |
|---|---|---|---|---|---|---|---|
| 1 | 10 | 4507445 | 4507493 | 49 | -4.05 | 39.5 | complex |
| 1 | 0 | 4510439 | 4510469 | 31 | -4 | 48.4 | no |
| 1 | 39 | 4510717 | 4510751 | 35 | -4.32 | 49.2 | no |
| 1 | 17 | 4520179 | 4520194 | 16 | -2.97 | 47 | no |
| 1 | 23 | 4529167 | 4529201 | 35 | -9.36 | 53.8 | no |
| 1 | 19 | 4538182 | 4538242 | 61 | -12.02 | 48.4 | no |
| 1 | 26 | 4539578 | 4539605 | 28 | -5.31 | 50.9 | no |
| 1 | 4 | 4603160 | 4603200 | 41 | -2.31 | 35.1 | MP |
| 1 | 0 | 4615841 | 4615896 | 56 | -5 | 44.6 | no |
| 2 | 22 | 2064544 | 2064558 | 15 | -2.18 | 46.2 | no |
| 2 | 0 | 2064783 | 2064812 | 30 | -3.94 | 51.7 | no |
| 2 | 0 | 2737152 | 2737176 | 25 | -4.6 | 51.4 | no |
| 3 | 24 | 2730177 | 2730272 | 96 | -13.96 | 41.8 | complex |
| 3 | 0 | 2732726 | 2732745 | 20 | -7 | 60.6 | no |
| 3 | 20 | 2733283 | 2733346 | 64 | -10.16 | 47.3 | no |
| 3 | 30 | 2739460 | 2739475 | 16 | -2.79 | 45.4 | no |
| 3 | 0 | 2739625 | 2739659 | 35 | -6.58 | 45.8 | extended |
| 3 | 24 | 2741098 | 2741151 | 54 | -10.82 | 50 | no |
| 3 | 24 | 2741752 | 2741805 | 54 | -10.42 | 45.1 | complex |
| 3 | 0 | 2762308 | 2762372 | 65 | -7.59 | 41.1 | no |
| 3 | 0 | 2762474 | 2762517 | 44 | -4.68 | 39.3 | no |
| 3 | 11 | 2787552 | 2787598 | 47 | -9.71 | 52.9 | no |
| 3 | 33 | 2789490 | 2789524 | 35 | -12.5 | 65.1 | no |
| 3 | 4 | 2790112 | 2790141 | 30 | -9.03 | 57.7 | no |
| 4 | 24 | 2818208 | 2818311 | 104 | -18.82 | 46.9 | no |
| 4 | 0 | 2818566 | 2818638 | 73 | -6.8 | 39.2 | no |
| 4 | 0 | 2865046 | 2865120 | 75 | -17.3 | 52.1 | no |
| 4 | 22 | 2864621 | 2864668 | 48 | -4.14 | 39.9 | no |
| 5 | 20 | 2406500 | 2406518 | 19 | -4.14 | 50.6 | no |
| 5 | 12 | 2432422 | 2432465 | 44 | -3.14 | 34 | no |
| 5 | 0 | 2447757 | 2447806 | 50 | -5.32 | 44.5 | extended |
| 5 | 0 | 2472736 | 2472753 | 18 | -5.08 | 60.8 | no |
| 5 | 0 | 2481869 | 2481901 | 33 | -9.61 | 65.8 | no |
| 5 | 4 | 2485778 | 2485812 | 35 | -9.49 | 58.7 | no |
| 6 | 22 | 2149229 | 2149258 | 30 | -5.79 | 43 | no |
| 6 | 0 | 2149500 | 2149539 | 40 | -3.7 | 41.7 | no |
| 6 | 27 | 2204457 | 2204473 | 17 | -2.56 | 56.2 | no |

| | | | | | | | |
|---|---|---|---|---|---|---|---|
| 7 | 15 | 2575496 | 2575522 | 27 | -7.17 | 54.3 | no |
| 7 | 6 | 2617694 | 2617735 | 42 | -5.59 | 41.5 | no |
| 7 | 18 | 2617863 | 2617903 | 41 | -5.21 | 44.2 | no |
| 7 | 0 | 2618350 | 2618368 | 19 | -2.65 | 44.1 | no |
| 7 | 20 | 2619032 | 2619048 | 17 | -3.33 | 45.9 | no |
| 7 | 0 | 2630477 | 2630522 | 46 | -8.83 | 48.6 | complex |
| 7 | 24 | 2603683 | 2603713 | 31 | -4.79 | 47.7 | no |
| 7 | 0 | 2636529 | 2636639 | 111 | -13.09 | 41.3 | no |
| 7 | 0 | 2646080 | 2646100 | 21 | -5.32 | 52.5 | no |
| 8 | 0 | 1814179 | 1814211 | 33 | -8.53 | 43.4 | MP |
| 8 | 19 | 1824932 | 1824951 | 20 | -3.3 | 47.5 | no |
| 8 | 0 | 1851716 | 1851740 | 25 | -3.18 | 45.5 | no |
| 8 | 37 | 1852408 | 1852432 | 25 | -7.13 | 58.4 | no |
| 8 | 21 | 1869439 | 1869466 | 28 | -3.05 | 45.5 | no |
| 8 | 0 | 1870116 | 1870139 | 24 | -12.28 | 79.7 | no |
| 8 | 0 | 1897027 | 1897077 | 51 | -5.33 | 42.8 | no |
| 8 | 25 | 1898595 | 1898622 | 28 | -5.12 | 56.1 | no |
| 8 | 22 | 1989215 | 1989252 | 38 | -4.97 | 46.3 | no |
| 8 | 24 | 1905265 | 1905349 | 85 | -10.56 | 39.8 | no |
| | | | | 40.68 | -6.8 | 48.5 | |

SI Table_3A. **Position and Stability of Palindromes Identified from the genomic DNA of centromeric TRP/INVs containing deletions of 40 bp or less.** The chromosomal (Chr) identity, deletion size, palindrome coordinates, palindrome length (stem + loop), and predicted ΔG and Tm values are listed together with the presence or absence of any extended or complex structures. The red color indicates palindromes assayed by direct sequencing and confirmed (and often extended) by an analysis of the surrounding region. The green hue indicates cases in which junctions were deduced from contiguous but discontinuous sequences. Red font indicates ΔG values above -3 kcal/mole.

| Chr | Deletion | Palindrome (start) | Location (stop) | Length (bp) | ΔG( kcal/mol) | Tm (°C) | Paired extensions |
|---|---|---|---|---|---|---|---|
| 1 | 1 | 475333 | 475351 | 19 | -2.76 | 53.6 | no |
| 1 | 16 | 873985 | 874034 | 50 | -6.61 | 44.4 | MP |
| 1 | 65 | 874682 | 874752 | 71 | -10.31 | 50.2 | MP |
| 1 | 19 | 937795 | 937823 | 29 | -5.41 | 61.9 | no |
| 1 | 26 | 938023 | 938060 | 38 | -2.32 | 37.5 | no |
| 1 | 0 | 2266091 | 2266116 | 26 | -9.28 | 68.8 | complex |
| 1 | 0 | 2286583 | 2286600 | 18 | -7.27 | 73.1 | no |
| 1 | 9 | 2300610 | 2300621 | 12 | -5.17 | 66.5 | no |
| 1 | 16 | 3240252 | 3240263 | 12 | -2.96 | 48.9 | no |
| 1 | 26 | 3436030 | 3436037 | 8 | -1.21 | 42.2 | no |
| 1 | 17 | 3436335 | 3436424 | 90 | -12.73 | 48 | complex |
| 1 | 11 | 3485742 | 3485768 | 27 | -8.89 | 60.7 | no |
| 1 | 3 | 3561732 | 3561750 | 19 | -2.42 | 39.6 | no |
| 1 | 23 | 3562955 | 3563012 | 58 | -8.2 | 44.2 | complex |
| 1 | 18 | 3750743 | 3750762 | 20 | -3.47 | 50.4 | no |
| 1 | 18 | 3751284 | 3751336 | 53 | -14.27 | 61.0 | no |
| 1 | 13 | 3893770 | 3893797 | 28 | -7.74 | 63.7 | extended |
| 1 | 7 | 4140360 | 4140412 | 53 | -6.91 | 45.2 | MP |
| 1 | 6 | 4140794 | 4140814 | 21 | -4.27 | 56.4 | no |
| 1 | 22 | 4459681 | 4459746 | 66 | -1.71 | 30.2 | no |
| 1 | 15 | 4648287 | 4648306 | 20 | -5.3 | 57.2 | no |
| 1 | 16 | 4648485 | 4648511 | 27 | -3.61 | 44.3 | no |
| 1 | 32 | 4868748 | 4868787 | 40 | -11.74 | 67 | no |
| 1 | 19 | 4988912 | 4988929 | 18 | -5.17 | 71.4 | no |
| 1 | 16 | 5016582 | 5016623 | 42 | -11.1 | 55.3 | no |
| 1 | 0 | 5016955 | 5016968 | 14 | -4.96 | 64.8 | no |
| 2 | 0 | 454491 | 454529 | 39 | -6.82 | 50.1 | complex |
| 2 | 0 | 454639 | 454669 | 31 | -6.86 | 57.4 | no |
| 2 | 0 | 493196 | 493210 | 15 | -4.72 | 68.6 | extended |
| 2 | 40 | 1804836 | 1804865 | 30 | -10.36 | 62.4 | no |
| 2 | 11 | 2305794 | 2305841 | 48 | -2.99 | 37.3 | complex |
| 2 | 20 | 2600637 | 2600646 | 10 | -3.63 | 52.6 | no |
| 2 | 30 | 2674415 | 2674485 | 71 | -5.72 | 41.5 | MP |
| 2 | 32 | 3246038 | 3246070 | 32 | -9.54 | 57 | no |
| 2 | 4 | 3348475 | 3348579 | 105 | -5.26 | 42.8 | complex |

| | | | | | | | |
|---|---|---|---|---|---|---|---|
| 2 | 12 | 3365066 | 3365266 | 201 | -5.53 | 78.5 | extended |
| 2 | 0 | 3365548 | 3365568 | 21 | -4.84 | 57 | MP |
| 2 | 0 | 3840022 | 3840121 | 100 | -10.34 | 42 | complex |
| 2 | 35 | 3840681 | 3840693 | 13 | -4.52 | 52.4 | no |
| 2 | 15 | 3894188 | 3894209 | 22 | -6.03 | 67.7 | no |
| 2 | 0 | 4116246 | 4116304 | 59 | -17.44 | 59.7 | no |
| 2 | 4 | 4171485 | 4171510 | 26 | -3.42 | 51.7 | extended |
| 2 | 0 | 4273901 | 4273927 | 27 | -7.05 | 57 | no |
| 2 | 0 | 5407963 | 5407985 | 23 | -13.94 | 83.9 | no |
| 2 | 40 | 5550965 | 5550988 | 24 | -3.98 | 53.5 | no |
| 2 | 17 | 5551234 | 5551263 | 30 | -9.01 | 70.6 | no |
| 3 | 8 | 268837 | 268854 | 18 | -3.07 | 49.7 | no |
| 3 | 13 | 269129 | 269150 | 22 | -4.75 | 60.1 | no |
| 3 | 0 | 567045 | 567060 | 16 | -4.75 | 61.1 | no |
| 3 | 0 | 573282 | 573312 | 31 | -8.35 | 69.9 | no |
| 3 | 0 | 616220 | 616250 | 31 | -9.18 | 59.7 | no |
| 3 | 27 | 961828 | 961857 | 30 | -4.43 | 45.2 | no |
| 3 | 7 | 1064407 | 1064426 | 20 | -3.87 | 51.2 | no |
| 3 | 21 | 1230017 | 1230054 | 38 | -8.02 | 54 | extended |
| 3 | 37 | 1230218 | 1230247 | 30 | -8.8 | 60.3 | no |
| 3 | 0 | 1526699 | 1526718 | 20 | -8.37 | 71.4 | no |
| 3 | 27 | 1572014 | 1572028 | 15 | -6.95 | 74.1 | extended |
| 3 | 0 | 1572702 | 1572727 | 26 | -9.18 | 68 | no |
| 3 | 0 | 1734350 | 1734376 | 27 | -6.42 | 50.8 | no |
| 3 | 18 | 2424655 | 2424677 | 23 | -4.56 | 44.7 | no |
| 3 | 11 | 2455727 | 2455754 | 28 | -3.67 | 55.3 | no |
| 3 | 17 | 2455926 | 2455945 | 20 | -2.47 | 43.4 | no |
| 3 | 21 | 2531330 | 2531360 | 31 | -3.22 | 48.6 | no |
| 3 | 23 | 2922514 | 2922535 | 22 | -8.24 | 72.6 | no |
| 3 | 30 | 2922806 | 2922830 | 25 | -8.76 | 68.1 | no |
| 3 | 20 | 3287357 | 3287368 | 12 | -2.85 | 53.2 | no |
| 3 | 0 | 3287741 | 3287757 | 17 | -4.78 | 56.9 | extended |
| 3 | 37 | 3293034 | 3293071 | 38 | -5.97 | 56.5 | no |
| 3 | 31 | 3837423 | 3837458 | 36 | -9.23 | 62.5 | extended |
| 3 | 25 | 3837616 | 3837637 | 22 | -5.56 | 72.3 | no |
| 3 | 0 | 3921549 | 3921567 | 19 | -1.84 | 40.5 | no |
| 3 | 24 | 3921753 | 3921774 | 22 | -7.72 | 65.8 | no |
| 3 | 34 | 4213341 | 4213382 | 42 | -6.32 | 49.9 | no |
| 3 | 16 | 4702849 | 4702860 | 12 | -2.36 | 47.4 | no |

| | | | | | | | |
|---|---|---|---|---|---|---|---|
| 3 | 25 | 4712272 | 4712306 | 35 | -10.54 | 77.3 | no |
| 3 | 25 | 4712627 | 4712716 | 90 | -7.02 | 39.3 | complex |
| 3 | 27 | 4939338 | 4939363 | 26 | -6.4 | 64.8 | extended |
| 3 | 0 | 4939733 | 4939746 | 14 | -2.24 | 45.1 | no |
| 3 | 2 | 4990641 | 4990657 | 17 | -2.99 | 53.7 | no |
| 3 | 37 | 4990857 | 4990878 | 22 | -6.4 | 66.6 | no |
| 4 | 7 | 43134 | 43152 | 19 | -7.92 | 69.6 | extended |
| 4 | 0 | 568414 | 568439 | 26 | -4.05 | 59.2 | extended |
| 4 | 7 | 568570 | 568593 | 24 | -5.55 | 62.4 | no |
| 4 | 8 | 694087 | 694103 | 17 | -4.45 | 62.1 | no |
| 4 | 0 | 794920 | 794944 | 25 | -9.95 | 70 | no |
| 4 | 30 | 870242 | 870261 | 20 | -3.28 | 65.6 | no |
| 4 | 2 | 1063138 | 1063169 | 32 | -6.05 | 62 | no |
| 4 | 10 | 1063422 | 1063444 | 23 | -5.01 | 56.6 | no |
| 4 | 0 | 1194263 | 1194283 | 21 | -6.12 | 69 | extended |
| 4 | 10 | 1194773 | 1194791 | 19 | -3.79 | 57.3 | no |
| 4 | 0 | 1273121 | 1273137 | 17 | -5.67 | 66.8 | no |
| 4 | 0 | 1274984 | 1275015 | 32 | -5.22 | 45.3 | no |
| 4 | 17 | 1275166 | 1275202 | 37 | -8.16 | 53.4 | no |
| 4 | 8 | 1607104 | 1607126 | 23 | 3.64 | 64.1 | no |
| 4 | 38 | 1607429 | 1607463 | 35 | -6.24 | 55.7 | no |
| 4 | 0 | 2528116 | 2528146 | 31 | -7.88 | 55.4 | no |
| 4 | 37 | 2567764 | 2567811 | 48 | -7.04 | 46.7 | complex |
| 4 | 40 | 3150093 | 3150134 | 42 | -5.88 | 51.4 | no |
| 4 | 25 | 3150305 | 3150395 | 91 | -12.43 | 43.9 | no |
| 4 | 8 | 3431397 | 3431420 | 24 | -8.63 | 69.1 | extended |
| 4 | 29 | 3869829 | 3869839 | 11 | -4.91 | 74.1 | no |
| 4 | 0 | 4238730 | 4238747 | 18 | -5.65 | 73.4 | no |
| 4 | 0 | 4239051 | 4239084 | 34 | -7.62 | 61.1 | no |
| 4 | 0 | 4240170 | 4240193 | 24 | -9.54 | 68.3 | no |
| 4 | 0 | 4414478 | 4414497 | 20 | -7.39 | 72.1 | extended |
| 4 | 16 | 4414743 | 4414764 | 22 | -4.91 | 57.8 | no |
| 4 | 17 | 4622841 | 4622861 | 21 | -8.95 | 69.5 | no |
| 5 | 33 | 345980 | 346025 | 46 | -8.97 | 51.5 | no |
| 5 | 24 | 346308 | 346323 | 16 | -2.94 | 59.9 | no |
| 5 | 5 | 375999 | 376059 | 61 | -6.52 | 41.5 | complex |
| 5 | 30 | 495783 | 495933 | 151 | -31.55 | 53.9 | no |
| 25 | 25 | 618600 | 618658 | 59 | -5.91 | 42.2 | no |
| 5 | 0 | 618917 | 618943 | 27 | -6.54 | 56.3 | extended |

| | | | | | | | |
|---|---|---|---|---|---|---|---|
| 5 | 17 | 636606 | 636616 | 11 | -2.71 | 54.8 | no |
| 5 | 8 | 1566416 | 1566465 | 50 | -11.39 | 55 | no |
| 5 | 14 | 1706110 | 1706136 | 27 | -7.36 | 72 | no |
| 5 | 39 | 1759115 | 1759167 | 53 | -6.95 | 43.8 | complex |
| 5 | 7 | 1760260 | 1760280 | 21 | -6.95 | 72.3 | no |
| 5 | 16 | 1847774 | 1847850 | 77 | -15.34 | 51.2 | no |
| 5 | 25 | 2017426 | 2017471 | 46 | -6.18 | 42.8 | no |
| 5 | 25 | 2017942 | 2017982 | 41 | -5.02 | 49.3 | no |
| 5 | 0 | 2331951 | 2331987 | 37 | -4.94 | 46.5 | extended |
| 5 | 0 | 3445422 | 3445437 | 16 | -4.6 | 61.5 | no |
| 5 | 0 | 3772422 | 3772439 | 18 | -7.07 | 71.7 | extended |
| 5 | 30 | 3772737 | 3772761 | 25 | -4.91 | 62.2 | no |
| 5 | 0 | 4111968 | 4111984 | 17 | -4.91 | 62.2 | no |
| 5 | 24 | 4257955 | 4257975 | 21 | -4.71 | 54.8 | no |
| 6 | 39 | 561314 | 561352 | 39 | -10.34 | 60.1 | complex |
| 6 | 11 | 561601 | 561684 | 84 | -14.51 | 48.6 | complex |
| 6 | 18 | 672921 | 672939 | 19 | -6.46 | 69 | no |
| 6 | 0 | 1167663 | 1167688 | 26 | -4.21 | 45.7 | no |
| 6 | 12 | 1167934 | 1167953 | 20 | -11.85 | 80.8 | no |
| 6 | 0 | 1661313 | 1661337 | 25 | -7.71 | 66.7 | extended |
| 6 | 25 | 1661471 | 1661508 | 38 | -7.26 | 54.7 | no |
| 6 | 0 | 2486550 | 2486570 | 21 | -12.11 | 87.5 | no |
| 6 | 0 | 2486789 | 2486806 | 18 | -4.13 | 62.2 | no |
| 6 | 0 | 2523428 | 2523443 | 16 | -4.68 | 57.6 | no |
| 6 | 21 | 2702045 | 2702094 | 50 | -9.92 | 56.9 | complex |
| 6 | 30 | 2745494 | 2745517 | 24 | -3.86 | 59 | no |
| 7 | 35 | 108999 | 109148 | 150 | -19.55 | 44.3 | no |
| 7 | 7 | 109686 | 109703 | 18 | -5.6 | 61.0 | no |
| 7 | 19 | 191281 | 191292 | 12 | 2.79 | 57.8 | no |
| 7 | 14 | 289804 | 289832 | 29 | -5.15 | 53.3 | no |
| 7 | 38 | 416292 | 416303 | 12 | -2.7 | 49.4 | no |
| 7 | 29 | 1169475 | 1169503 | 29 | -4.85 | 51.5 | no |
| 7 | 0 | 1656247 | 1656296 | 50 | -12.75 | 53.2 | no |
| 7 | 16 | 1765307 | 1765334 | 28 | -3.9 | 50.5 | no |
| 7 | 0 | 1814640 | 1814658 | 19 | -4.36 | 67.9 | extended |
| 7 | 18 | 1864743 | 1864755 | 13 | -3.03 | 52.1 | no |
| 7 | 22 | 1865242 | 1865272 | 31 | -6.85 | 55.3 | no |
| 7 | 26 | 2298835 | 2298853 | 19 | -4.77 | -4.77 | no |
| 8 | 0 | 256532 | 256552 | 21 | -4.3 | 52 | extended |

| | | | | | | | |
|---|---|---|---|---|---|---|---|
| 8 | 28 | 731310 | 731346 | 37 | -7.93 | 58.6 | complex |
| 8 | 17 | 797320 | 797339 | 20 | -7.46 | 75.4 | no |
| 8 | 27 | 809132 | 809157 | 26 | -6.66 | 55.8 | no |
| 8 | 6 | 809332 | 809357 | 26 | -6.67 | 66.3 | no |
| 8 | 10 | 841466 | 841482 | 17 | -2.41 | 47.9 | no |
| 8 | 20 | 919877 | 919917 | 41 | -6.26 | 45.2 | no |
| 8 | 31 | 1428024 | 1428055 | 32 | -6.88 | 55 | no |
| 8 | 0 | 1428359 | 1428392 | 34 | -7.82 | 63.6 | no |
| 8 | 7 | 1769614 | 1769642 | 29 | -9.79 | 69.9 | no |
| 8 | 11 | 1769919 | 1769934 | 16 | -3.28 | 58 | no |
| 8 | 37 | 2160569 | 2160612 | 44 | -7.14 | 57 | complex |
| 8 | 6 | 2314475 | 2314490 | 16 | -5.06 | 62.3 | extended |
| 8 | 5 | 2319380 | 2319434 | 55 | -11.6 | 54.8 | extended |
| 8 | 2 | 2738233 | 2738252 | 20 | -6.29 | 64.6 | no |
| 8 | 22 | 2738528 | 2738548 | 21 | -4.64 | 59 | no |
| 8 | 21 | 2804431 | 2804481 | 51 | -3.99 | 37.4 | no |
| 168 | | | | 33.2 | -6.62 | 56.4 | |

SI Table_3B. **Position and Stability of Palindromes Identified from the genomic DNA of dispersed TRP/INVs containing deletions of 40 bp or less.** Samples were assayed as described in SI Tabe 3A. Multipalindromic structures (MP) are also shown.

| Chr | Deletion | Palindrome (start) | Location (stop) | Length (bp) | ΔG ( kcal/mol) | Tm (°C) | Paired Extensions |
|---|---|---|---|---|---|---|---|
| 1 | 37 | 13359 | 13394 | 36 | -7.94 | 51.6 | extended |
| 1 | 0 | 75341 | 75397 | 57 | -5.76 | 38.3 | no |
| 1 | 24 | 77036 | 77067 | 32 | -4.15 | 46.2 | extended |
| 2 | 26 | 409658 | 409705 | 48 | -12.39 | 54.3 | no |
| 2 | 23 | 3673359 | 3673405 | 47 | -8.48 | 49.8 | no |
| 3 | 0 | 2692978 | 2692999 | 22 | -3.22 | 44.3 | no |
| 3 | 0 | 2705424 | 2705444 | 21 | -2.99 | 44.2 | no |
| 3 | 32 | 5171545 | 5171569 | 25 | -4.5 | 58.3 | extended |
| 3 | 32 | 5191849 | 5191894 | 46 | -7.53 | 48.2 | no |
| 4 | 40 | 2071 | 2111 | 41 | -3.98 | 44.3 | extended |
| 4 | 0 | 3583 | 3602 | 20 | -3.88 | 45.2 | no |
| 4 | 36 | 5539 | 5569 | 31 | -4.33 | 60.2 | extended |
| 5 | 0 | 969 | 1028 | 60 | -7.92 | 42.7 | no |
| 5 | 0 | 1720 | 1740 | 21 | -4.88 | 52.1 | no |
| 5 | 0 | 2201 | 2224 | 24 | -2.22 | 38.8 | no |
| 5 | 27 | 3808 | 3906 | 99 | -8.83 | 45.1 | no |
| 5 | 12 | 4511776 | 4511811 | 36 | -6.77 | 55.4 | no |
| 5 | 0 | 4510716 | 4510736 | 21 | -1.21 | 34 | no |
| 7 | 0 | 690480 | 690566 | 87 | -13.45 | 44 | no |
| 7 | 17 | 690286 | 690319 | 34 | -6.66 | 48.7 | no |
| 8 | 20 | 6022 | 6039 | 18 | -2.67 | 46.8 | no |
| 8 | 0 | 3238765 | 3238854 | 90 | -18.55 | 51.7 | no |
| 8 | 37 | 3239037 | 3239114 | 78 | -11.24 | 45.9 | no |
| 8 | 36 | 3241327 | 3241353 | 27 | -7.96 | 59.6 | no |
| 24 | | | | 42.5 | -6.73 | 47.9 | |

SI Table_3C**. Position and Stability of Palindromes Identified from the genomic DNA of subtelomeric TRP/INVs containing deletions of 40 bp or less**. Samples were assayed as described in SI Tabe 3A.
